# Parameter estimation in the age of degeneracy and unidentifiability

**DOI:** 10.1101/2021.11.28.470243

**Authors:** Dylan Lederman, Raghav Patel, Omar Itani, Horacio G. Rotstein

**Affiliations:** Federated Department of Biological Sciences, New Jersey Institute of Technology and Rutgers University; Department of Computer Sciences New Jersey Institute of Technology

## Abstract

Parameter estimation from observable or experimental data is a crucial stage in any modeling study. Identifiability refers to one’s ability to uniquely estimate the model parameters from the available data. Structural unidentifiability in dynamic models, the opposite of identifiability, is associated with the notion of degeneracy where multiple parameter sets produce the same pattern. Therefore, the inverse function of determining the model parameters from the data is not well defined. Degeneracy is not only a mathematical property of models, but it has also been reported in biological experiments. Classical studies on structural unidentifiability focused on the notion that one can at most identify combinations of unidentifiable model parameters. We have identified a different type of structural degeneracy/unidentifiability present in a family of models, which we refer to as the Lambda-Omega (Λ-Ω) models. These are an extension of the classical lambda-omega (*λ*-*ω*) models that have been used to model biological systems, and display a richer dynamic behavior and waveforms that range from sinusoidal to square-wave to spike-like. We show that the Λ-Ω models feature infinitely many parameter sets that produce identical stable oscillations, except possible for a phase-shift (reflecting the initial phase). These degenerate parameters are not identifiable combinations of unidentifiable parameters as is the case in structural degeneracy. In fact, reducing the number of model parameters in the Λ-Ω models is minimal in the sense that each one controls a different aspect of the model dynamics and the dynamic complexity of the system would be reduced by reducing the number of parameters. We argue that the family of Λ-Ω models serves as a framework for the systematic investigation of degeneracy and identifiability in dynamic models and for the investigation of the interplay between structural and other forms of unidentifiability resulting on the lack of information from the experimental/observational data.

## 1 Introduction

Mathematical models are useful tools that can be used to interpret experimental or observational data, make quantitative predictions that are amenable for experimental testing, and identify the mechanisms that underlie the generation of pattern of activity in terms of the interactions among the system’s components [1–3]. An important step in connecting models with experimental or observational data is to estimate the model parameters by fitting the model outputs to the available data (Fig. 1). The ability of models to make predictions, provide mechanistic explanations, and be useful for decision making all depends on the accuracy and reliability of the parameter estimation process. A large number of parameter estimation tools are available to scientists as well as methods to discover data-drive nonlinear dynamic equations [4–20] (and references therein) and tools to link data with models continue to develop.

**Figure 1:**
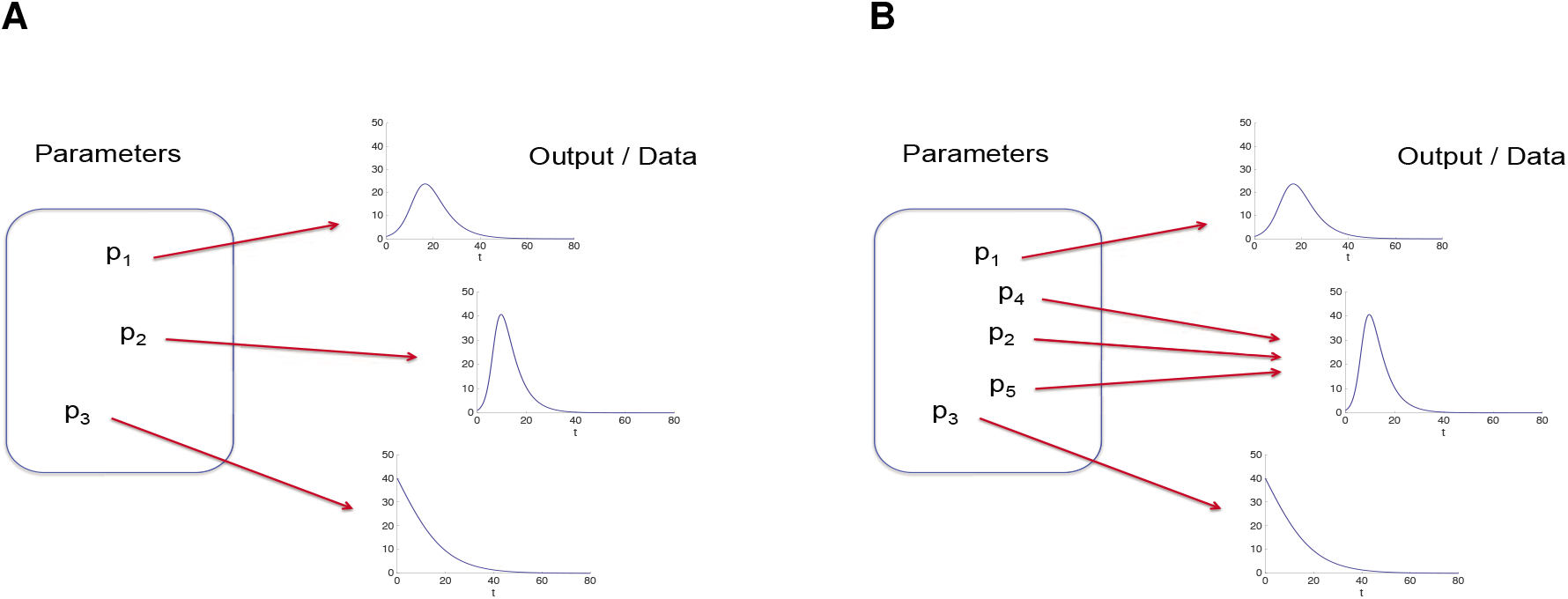
Representative parameter estimation diagram. The model parameter sets are represented by *p*_1_, *p*_2_, *p*_3_, *p*_4_ and *p*_5_ (within the framework on the left of each panel). The data (or data sets), either experimental/observational data or ground truth (output) model data, are represented by the graphs on the right of each panel. **A.** Schematic example of identifiability. Each data set is generated by one and only one parameter set. **B.** Schematic example of unidentifiability. Some data sets are generated by more than one parameter set.

A key feature of these tools is the minimization of an error function, a metric of the difference between the model output (simulations using estimated parameters) and the available (target) experimental/observed data. These data can be continuous (data collected with a high-frequency sampling, as compared to the scale of the process of interest, so that one can make the continuous approximation for mathematical purposes) or discrete (point process capturing the occurrence of events of interest). Error functions can be constructed by using all the data available in a point-to-point fashion (e.g., Fig. 2-A) or by computing one or more *attributes* that characterize these data (e.g., oscillation frequency, oscillation amplitude, oscillation duty cycle, stationary states, characteristic raise and decay times; Fig. 2-B).

**Figure 2:**
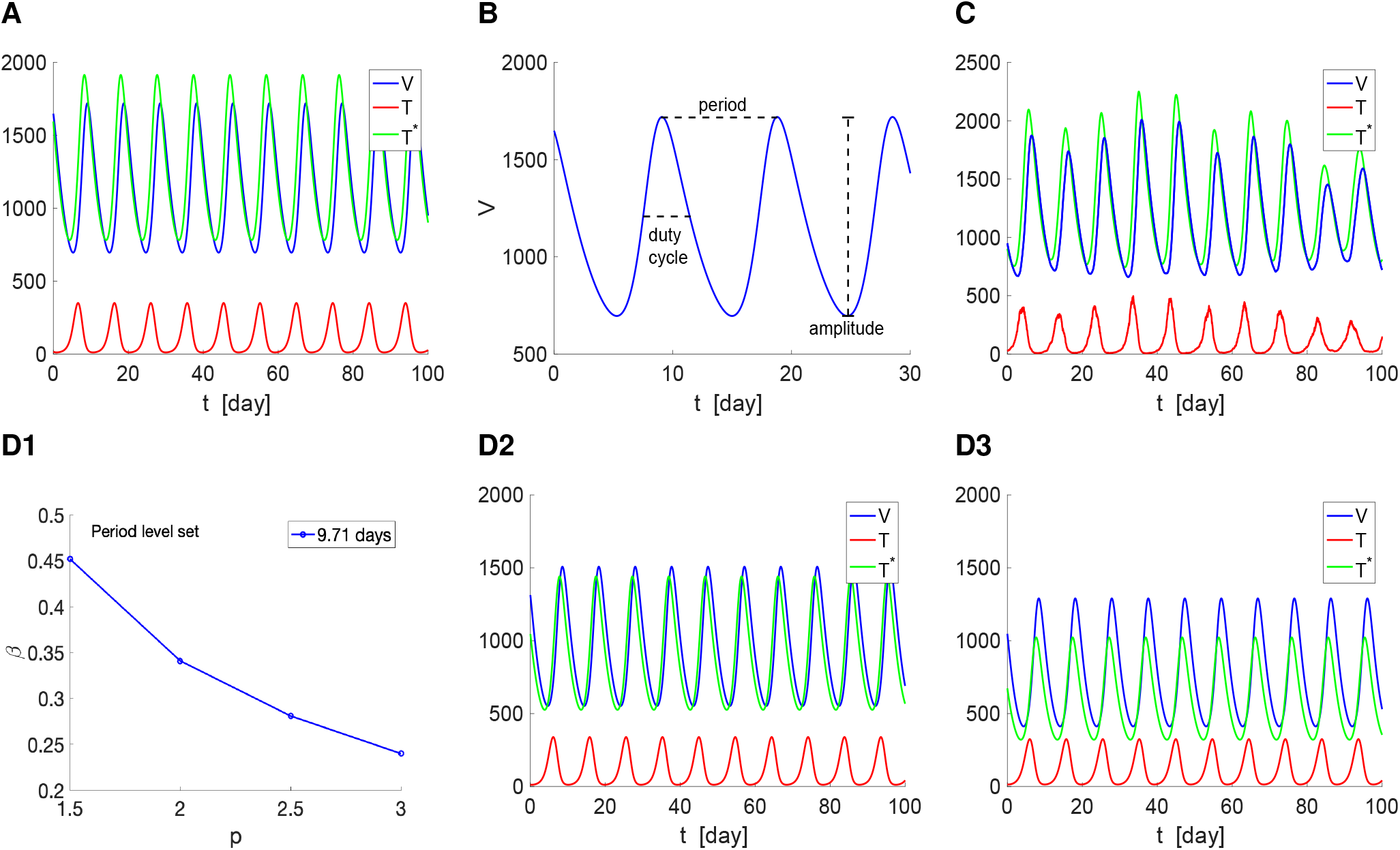
Dynamics and attributes for the oscillatory patterns in a HIV-1 model. The variables represent the concentrations of uninfected cells (*T*), infected cells (*T* *) and virus particles (*V*). **A.** Representative oscillatory pattern. The oscillation period is ~ 9.71 days. **B.** Three attributes for *V*: period, amplitude and duty cycle. These attributes can be computed similarly for *T* and *T* *. These attributes can be extracted from data if the data is regular enough. **C.** Representative oscillatory pattern for a stochastic logistic growth rate *p* of the T-cells. The mean period is ~ 9.82 days. The mean period after 10 trials is ~ 9.61 days. The average frequency of the irregular pattern in Fig. 2-C is almost equal to the frequency of the regular pattern in Fig. 2-A. Both were created using the same parameter values, except for the rate of logistic growth of the uninfected T-cells, which was constant in Fig. 2-A and normally distributed around this constant value in Fig. 2-C. Therefore, the frequency can be used as a reliable attribute, but not necessarily the amplitude, whose quantification may require statistical processing. We used the HIV-1 model (1)–(3) [21] (see also [22, 23]) with the following paramter values: *δ* = 10, *α* = 0.02, *p* = 3, *β* = 0.24, *Tmax* = 1500, *γ* = 2.4, *k* = 0.0027, *N* = 10 and *i* = 1. In panel C, *p* was drawn from a normal distribution with mean zero and and variance equal to 2. We rejected the negative values of *p* (the negative values of *p* were substituted by zero). **D.** Period level set in the *p*-*β* parameter space for the same period as in panel A (~ 9.71 days).

There are several difficulties associated with the implementation of parameter estimation tools. These difficulties are data-related (lack of access to all state variables, inconsistent gaps across trials), computational (algorithmic nature), statistical (data is noisy and therefore one can at best expect to estimate distributions of parameter values around a “true" mean), and structural (degeneracy, mathematical nature).

In an ideal situation, there would be a unique parameter set that fits the experimental/observational data (e.g., Fig. 1-A). We refer to the underlying model as *identifiable* [24]. In mathematical terms, the function linking parameter sets and data is bijective. In practice, these systems are subject to fluctuations from uncertain sources and one obtains a distribution of parameter sets (in a high-dimensional spaces whose dimensions are equal to the number of parameters of interest) centered at the “true" parameter set. The former has been termed *structural identifiability*, while the second has been termed *practical identifiability* or *estimability* [25]. Roughly speaking, the narrower the estimated parameters distribution (the smaller the variance), the more precise the parameter estimation process. However, this assumes that the center of the distribution is a good approximation to the “true" value. When necessary, the necessary validation of this assumption and the circumstance under which it is true can be done by using ground truth data generated by mathematical models. When continuous data is too noisy to be amenable for parameter estimation purposes, one might be able to extract useful discrete data in the form of attributes that capture the most relevant aspects of these data or point processes that are embedded in the data (e.g., Fig. 2-C and Fig. 3). However, this is done at the expense of loosing information and therefore has consequences for the accuracy of the parameter estimation process. Additional issues emerge when not all state variables are accessible or if gaps in discrete data sets are inconsistent across trials, requiring the use of imputation tools.

**Figure 3:**
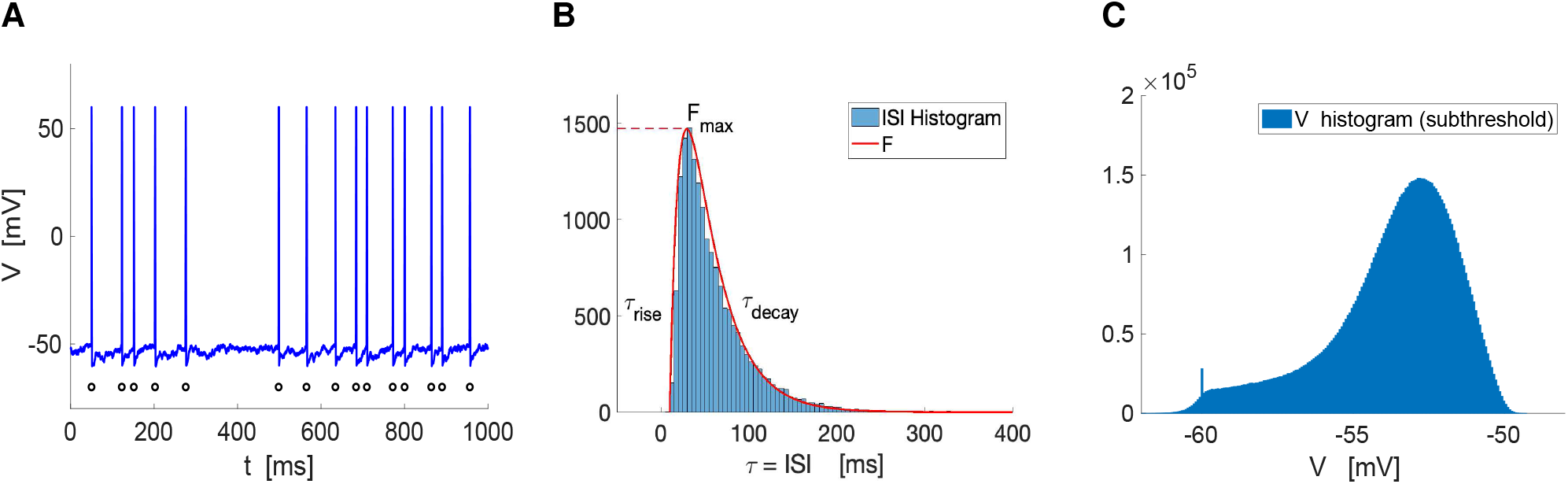
Dynamics and attributes of a representative neuronal spiking pattern. **A.** Membrane potential *V* (blue) and spike times (black). The attributes are the *V* trace (curve of *V* as a function of *t*) or the spike times sequence if one has no access to *V* during the interspike intervals (ISIs) or chooses not to use that information (e.g., if it is too noisy). **B.** Interspike interval (ISI) histogram (light blue) and approximation using an difference of exponentials *F* (red). The ISI sequence is defined as the difference of two consecutive spike times. *F* is defined by 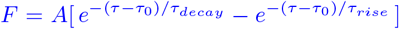. The attributes are *F_max_*, *τ_decay_*, *τ_rise_* and *τ*_0_. **C.** Subthreshold membrane potential *V* histogram. The peak at *V* = −60 corresponds to the transition from spiking to the reset voltage *V_thr_*. The mean is equal to −53.71 and the variance is equal to 4.63. We used a leaky integrate-and-fire (LIF) model in the noise-driven regime. The model is defined by eq. (59), describing the dynamics of the so-called passive cell (an RC circuit) with the addition of a voltage threshold *V_thr_* for spike generation and a reset values *V_rst_* after a spike has occurred. Spikes (action potentials) are generated when *V* reaches *V_trh_*, after which *V* resets to the prescribed value *V_rst_* (below the so-called resting potential or stable equilibrium of the underlying circuit). In the absence of noise, *V* reaches an equilibrium and therefore spikes are created by the presence of noise. For higher values of *Iapp* (and no noise), *V* spikes periodically. When noise is added, it disrupts the periodicity of the oscillations. Even if one has access to the subthreshold *V* data, the error minimization process might be not easy to accomplish because of the noisy subthreshold behavior. If this is the case, one can use the spike-times (panel A, black dots), a discrete variable, at the expense of losing some information, and the spike-time statistics (panel B) to characterize the data. We used the following parameter values *C* = 1 *μ* F/cm^2^, *G_L_* = 0.1 mS/cm^2^, *I_app_* = 0.8 mA/cm^2^, *E_L_* = 60 mV, *V_thr_* = −50 mV, *Vrst* = −60 mV and a white noise variance *D* = 0.25. We run the simulation for a total time of 1,000,000 ms with a time step Δ*t* = 0.1 ms.

*Unidentifiability*, the opposite of identifiability [25–31], is associated with the concept of degeneracy. Degeneracy refers to these situations where multiple sets of parameter values can produce the same observable output (e.g.,Fig. 1-B, Fig. 2-D1 if the output is the period), therefore making the inverse problem (of finding parameters given the data) ill-posed. In mathematical terms, the function linking parameter sets and data is not injective (at most surjective). (There is in fact an implicit assumption that these links are surjective.) Degeneracy is not a problem associated with the statistical uncertainty in the knowledge of a unique parameter set from which the data is generated referred to above, but a structural problem inherent to mathematical models where the same patterns (e.g., temporal, spatial) can be obtained from multiple parameter sets. Degeneracy gives rise to the concept of *level sets* in parameter space (e.g., Fig. 2-D1) [32, 33]. These are geometric objects (curves, surfaces, hypersurfaces) along which a given attribute of activity remains constant (see additional examples and a detailed discussion in the Appendix A), and are the geometric instantiation of the idea that a number of well-defined combinations of unidentifiable parameters can form an identifiable set. In other words, one can at most identify combinations of unidentifiable parameters.

Identifiability and unidentifiability have been studied in a number of systems. Structural identifiability has been studied using different approaches, including differential algebra [27, 34–39], Taylor series [40], similarity transformations [41, 42] and the Fisher information matrix [26, 43–45] (see also [46–48]). In recent work [32] we have used numerical simulations and dynamical systems tools to characterize the frequency and duty cycle level sets and explain what are the dynamic mechanisms responsible for their generation in the FitzHugh-Nagumo (FHN) [49] model and the Morris-Lecar model [50, 51] in the oscillatory regimes. Within a given attribute level set (e.g., period), the oscillatory patterns are non-identical (similarly to Fig. 2-D), and therefore dynamic balances operate to maintain the attribute of interest. The Morris-Lecar model is a neuronal model describing the interplay of two ionic currents (calcium and potassium), a leak (linear) current and a capacitive current. The FHN model is a caricature model that has been used to capture the oscillatory behavior in a number of fields (see Appendix A.3). In the relaxation oscillations regime, the oscillatory behavior of the FHN model is qualitatively similar to the one exhibited by the Morris-Lecar model and other models of biological relevance [2, 52, 53]. When the time scales are not well separated, the FHN model exhibits oscillations qualitatively similar to the HIV model in Fig. 2.

The use of non-trivial attributes makes the comparison between data and model output computationally less expensive; instead of having to use a large set of points (e.g., the histogram envelope captured by the red curve in Fig. 3-B or the full time courses in Fig. 2) one can use a significantly smaller number of attributes. In general, attributes refer to the parameters that characterize the data according to the way one chooses the data to be organized. On one extreme, the attributes are the whole set of data points and one ideally has access to all the model’s state variables. On the other extreme, the attributes are the minimal number of “numbers" necessary to fully characterize the available data. For example, frequency and amplitude are enough to describe a sinusoidal oscillation. A third attribute would be necessary if this oscillation is not centered at zero. Additional attributes such as the time constant governing the transition from an active (up) to a passive (down) phases would be necessary to discriminate between sinusoidal and relaxation oscillations. In the example in Fig. 2 the oscillations can be characterized by the frequency, amplitude and duty cycle. In the example in Fig. 3, the data consists of the interspike (or interpeak) intervals (ISIs or IPIs) histogram, which releases us from making detailed spike-time comparisons that could be strongly affected by noise and therefore complicates the error minimization process. This histogram can be characterized by four attributes (Fig. 3-B): the histogram peak *F_max_*, the characteristic rise and decay constants *τ_rise_* and *τ_decay_*, respectively, and the value of *τ* at which the histogram “begins", *τ*_0_. Additional attributes are the mean and variance of the ISI distribution. (Neuronal spiking behavior typically approximates a Poisson process [54, 55] and therefore the mean ISI is equal to its standard deviation.) Similar type of histograms could be computed for the frequency, amplitude and duty cycle in the HIV-1 viral kinetics example in Fig. 2-C. Note that attributes are used in connection with the data, while parameters are used in connection to the model used to generate or describe these data. A detailed discussion about the relationship between attributes (of patterns of activity) and model parameters in simple models including the prototypical FitzhHugh-Nagumo oscillator [49] and a Hepatitis C viral model [56] (see also [57]) is presented in the Appendix A.

Oscillations are ubiquitous in dynamical systems, particularly in biological systems [2,21,52,53,58–63] where degeneracy has been observed, not only in models, but also in experiments [64–66] (see also [32, 33]). The mechanisms underlying the presence of degeneracies in biological oscillations are not well understood. It is also unclear how the presence of degeneracies affect the parameter estimation process for models describing biological oscillations and oscillations in general.

We have identified a family of canonical models for unidentifiability/degeneracy where the limit cycle trajectories are unidentifiable and the degeneracy can be analytically computed. There are infinitely many parameter sets that produce identical stable oscillations, except possible for a for a phase-shift, which depends on the initial phase. In other words, all the attributes of the of the oscillatory patterns are degenerate and have the same level sets, which can be analytically computed. We refer to the generic formulation of this model as the Λ-Ω model. It is a generalization of the so-called *λ*-*ω* system [67, 68], also referred to as the Poincaré oscillator [69, 70]. These are real-valued special cases of the complex Ginzburg-Landau equation [71–74] and their role for modeling biological systems are discussed in detail in [58] (see references therein). The general formulation can exhibit bistability and a number of bifurcations. The classical Λ-Ω exhibit sinusoidal oscillations. Extended versions of this model produce oscillations with more complex waveforms, including relaxation and spike-like oscillations, and a rich repertoire of dynamic behavior, including bistability and even multistability. In contrast to other models showing structural unidentifiability due to redundancy in the number of parameters with physical meaning (see Appendix A for a detailed discussion), the number of parameters in the Λ-Ω models cannot be reduced without reducing the model complexity. To our knowledge, the issues of unidentifiability and degeneracy in the family of Λ-Ω models has not been raised before.

Because of the properties described above, the family of Λ-Ω models are an ideal system to ask fundamental questions about degeneracy of dynamical systems and their relationship with parameter estimation unidentifiability. The goal of this paper is to address these issues. Our results contribute to the development of a framework for the investigation of unidentifiability in dynamic models.

## 2 Methods

### 2.1 A viral infection kinetic model

For illustrative purposes we will use the HIV viral infection model presented in [21] (see also [22, 23]), which, as noted in [23] belongs to a more general class of viral infection models [75]. The model is described by the following equations for the concentration *T* of the uninfected T-cells, the concentration *T* * of productively infected T-cells, and the concentration *V* of free virus particles in the blood

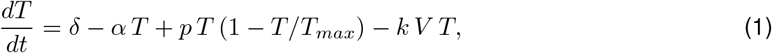

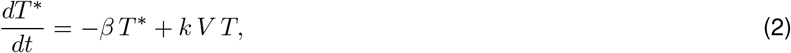

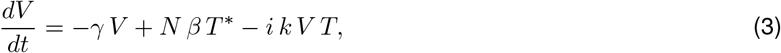

where *α* (day^−1^), *β* (day^−1^) and *γ* (day^−1^) are the death rates of the uninfected T-cells, the infected T-cells and the virus particles, respectively, *k* (mm^3^ day^−1^)is the contact rate of the uninfected T-cells and the viral particles, *δ* (mm^−3^ day^−1^) represents the constant production of the T-cells, *N* is the average number of virus particles produced by an infected T-cell, *p* (day^−1^) and *T_max_* (mm^−3^) are the growth rate and the carrying capacity, respectively, associated to the logistic growth of uninfected T-cells in the absence of virus particles, infected T-cells and other factors. As shown in Fig. 2, this model displays oscillatory behavior. Oscillations are typically observed in a number of virus dynamical systems [21, 76–78].

### 2.2 Lambda-Omega (Λ-Ω) models: general formulation

The Λ-Ω systems have the general form

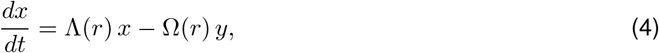

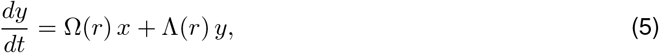

with

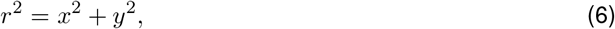

where *x* and *y* are state variables and *t* is time. The functions Λ and Ω are typically chosen to be polynomials of degree two or four [67,69,70], but they could in principle be well-behaved enough functions with the necessary properties to support oscillations as discussed in this paper.

### 2.3 Modified Λ-Ω models: general formulation

The modified Λ-Ω modes have the general form

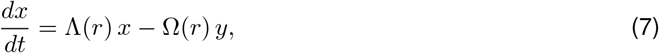

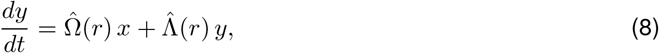

where the functions 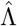 and 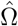 are different from the functions Λ and Ω, but belong to the same class. We use this formulation to investigate the consequences of the break of symmetry present when 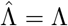 and 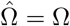.

### 2.4 Parameter estimation algorithms

To study the impacts of the Λ-Ω model’s degenerate mathematical structure on the estimation of its parameters, the behavior of three parameter estimation algorithms, covering a range of methods, were characterized. These include a Genetic Algorithm (GA) [10,11,79,80], Gradient Descent Algorithm [6,81], and Sequential Neural Posterior Estimation (SNPE) method [15, 82, 83]. The purpose of each of them is to find parameter values that fully minimize the difference between estimated solutions produced by the algorithm’s output and a data set representing the target (optimal) solution to the model. In practical applications of parameter estimation, experimental (or observational) data typically acts as this set of target data, however, for the purpose of this study, when working with the *λ*-*ω* model a simulated output using a predetermined set of parameters (*λ* = 1, *ω* = 1, *a* = 1, *b* = 1) acts as a ground truth data set (GTD) to stand in as a proxy for experimental data. To categorize the impacts of noise on the identifiability of the model’s parameters, each parameter estimation algorithm was tested using data sets with variable levels of noise applied to the GTD. We used Gaussian white noise with mean zero and variance *D*(*D* = 0, 100, 250), added to the both equations in the *λ*-*ω* model. This approach seeks to activate the transient dynamics of the system by perturbing its trajectory from the limit cycle. Under certain conditions, the activation of transients is believed to resolve some types of unidentifiability issues. Specifically in the case of the *λ*-*ω* model, if one is able to estimate *λ* from the transient dynamics, then the estimation of b is straightforward, eliminating the unidentifiability of the two parameters. In our simulations we used Δ*t* = 0.01.

#### 2.4.1 Stochastic Optimization: Genetic algorithms (GA)

The GA is an evolutionary algorithm based on the principles of natural selection, in which only “fit" individuals of a given generation are used to create subsequent generations [79, 80] (see also [10–13]). The fitness of parameter sets within the population are those that produce an output most closely resembling the GTD. In our numerical experiments, we quantify this as the difference between the ground truth and the simulated differences and amplitudes. However, other implementations of the algorithm can consider the attributes independently or simply use the time series produced by each parameter set as a whole. Both of these approaches were tested and showed no improvement on the identifiability of parameters. We used a multiobjective GA, where the estimation process begins by creating an initial population of randomly distributed parameter sets [79]; for the purposes of this study, populations were created using random values within the range of [0, 2]. In each generation, only the top half of the population that most optimally matches the GTD are used as “parents" to build the next generation. The next generation consists of both crosses between two independent parent sets and the original parent sets themselves with random mutations being added to parameter values at a rate of 1 over the number of parameters estimated. The number of generations produced and population size is specified by the user, with executions in our study using 500 generations and populations of 250 individuals. To ensure the adequacy of these conditions, trials with a larger population size (up to 2000 individuals) and/or higher number of generations (up to 1000 generations) were tested and shown to have no impact on the performance of the GA.

#### 2.4.2 Sequential Neural Posterior Estimation (SNPE)

We implemented SNPE using the simulation-based inference (SBI) developer toolbox [15, 82, 83]. SNPE falls under a larger category of algorithms called Sequential Neural Processes, which use Bayesian inference to estimate parameter values [15, 82, 83]. This technique is a statistical method for calculating likelihood of events based on prior observations, and it is applied to determine which parameter values are most likely to produce the behavior observed in the GTD [15, 82, 83]. Similarly to the GA, Sequential Neural Processes start with an initial distribution of parameters, which is then refined, in this case using Bayesian inference. For this application of SNPE, an initial uniform distribution on [0, 2] is created as a prior distribution. Then, 10000 simulations of the model are created by sampling from parameter sets within this distribution in order to infer a posterior distribution based on which sets produce an output most closely matching the GTD.

## 3 Results

### 3.1 Unidentifiability and degeneracy: a preliminary discussion using simple models

Here we introduce some ideas related to unidentifiability and degeneracy in dynamical systems, including the concepts of activity attributes, level sets and parameter redundancy in models and their outputs. We do this in the context of relatively simple models. A detailed mathematical description of these models and the unidentifiability analysis are presented in the Appendix A.

In the 1D minimal model example discussed in the Appendix A.1, the minimal model has no level sets. The number of minimal model parameters is equal to the number of attributes that characterize the pattern. In fact, the minimal model parameters are themselves the attributes. In the 2D minimal model example discussed in the Appendix A.2, there are no level sets for the time constant *τ* and this attribute is determined uniquely by the minimal model parameter *b* in eq. (63). In contrast, the damped oscillations frequency *ω* (64) is a combination of the minimal model parameters *b* and *c* and has level sets in this parameter space. Clearly, if one can estimate the two attributes *τ* and *ω*, one can identify the minimal model parameters *b* and *c*. If one can only estimate *ω*, then the model has degeneracy. Typically, by design, the minimal model parameters have no physical (or biological) meaning, but only dynamic meaning. They are a minimal set of model parameters in that eliminating one of them reduces the dynamic complexity.

The situation changes when models are created to reproduce or explain experimental or observational data. The number of physically meaningful model parameters is larger than the number of minimal model parameters needed to maintain the model dynamic complexity. The level sets and degeneracy that we find in the realistic models in both cases (Appendices A.1 and A.2) emerge because the minimal model parameters (with dynamic meaning) are combinations of (realistic) parameters with physical meaning. From that perspective, the realistic parameters generate a dynamic redundancy, which is reflected by the fact that the minimal model parameters are combinations of physical parameters. This type of degeneracy cannot be resolved unless one has access to additional data.

The FHN model discussed in the Appendix A.3 is a nonlinear minimal model in the oscillatory regime. In the example presented in the Appendix A.3, there are level sets, which emerge as two or more minimal model parameter sets dynamically balance each other to maintain an attribute constant (e.g., frequency, see Fig. 17-A and -B) [32] similarly to the HIV model oscillations discussed in the context of Fig. 2. Attribute level sets such as frequency, amplitude, and duty-cycle do not coincide except perhaps in some specific cases [32]. Additional degeneracies emerge in realistic models (e.g., Morris-Lecar model [32]) as the number of parameters with biological meaning increases with the consequent increase in parameter redundancies. However, it is not always possible to reduce nonlinear models to minimal models.

Fig. 17-C in the Appendix A.1 illustrates the fact that additional attributes may be necessary to characterize oscillatory patterns. The nonidentical sinusoidal and square-wave functions have identical frequency, amplitude and duty cycle. This raises the need of identifying the appropriate attributes to characterize a given pattern and their optimal number.

### 3.2 The Λ-Ω systems are canonical models for unidentifiability/degeneracy

Here we discuss the role of the Λ-Ω models generically described by eqs. (4)–(6) as canonical models for unidentifiability where a limit cycle with the same frequency can be generated by multiple combinations of model parameters.

A change of coordinates from Cartesian to polar

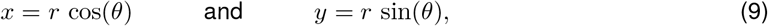

transforms system (4)–(5) into

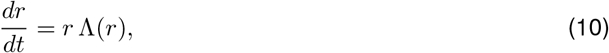

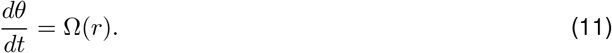

The equation for the radius of the limit cycle is independent of the the angular velocity *θ*. Therefore, the limit cycles are circles around zero and their radii 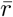 are the solutions of

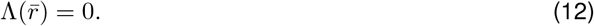

The number of limit cycles 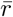 depends on the form and complexity of the function Λ(*r*). Since *r >* 0, a limit cycle of radius 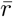 is stable if and only if 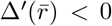. The frequency of the solution along this limit cycle is 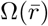 and the period is 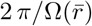. Because, if a stable limit cycle exists, in the limit of *t* → ∞ the dynamics are described completely by 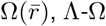 systems are often referred to as phase oscillators. Note that 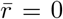 corresponds to a fixed-point and therefore the Λ-Ω systems always have a fixed-point whose stability properties also depend on the sign of Δ^′^(0).

Limit cycle and frequency degeneracy emerges when the function Δ is defined by two or more parameters. Additional degeneracy in the frequency results from the parameters defining the function Ω.

### 3.3 Lambda-omega systems of order 2: limit cycle degeneracy

Λ-Ω systems of order 2 (Λ-Ω_2_) correspond to the following quadratic choices for Λ(*r*) and Ω(*r*)

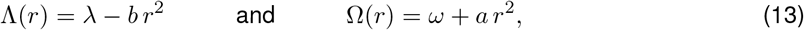

where *ω*, *λ*, *a* and *b* are constant. Eqs. (4)–(5) become

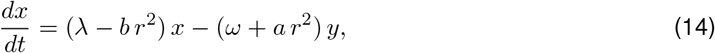

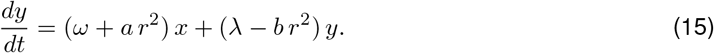

Following [67], we refer to these models as *λ*-*ω* models. They have been also referred to as Poincaré oscillators [69] and are real -valued special cases of the complex Ginzburg-Landau equation as mentioned above [71–74].

The parameters *λ* and *ω* control the linear dynamics and the parameters *a* and *b* control the contribution of the nonlinear terms to the system’s dynamics. In this sense, the *λ*-*ω* model is minimal. As we will see, the parameters *λ* and *ω* play a stronger role in controlling the transient dynamics than the parameters *a* and *b*.

#### Limit cycles, fixed-points, stability and oscillation frequency

These *λ*-*ω* systems have a single limit cycle for

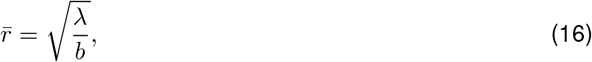

if *λ/b >* 0, which is stable if and only if *λ <* 0. Otherwise, the fixed-point 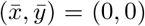 is stable. Along the limit cycle, the frequency is constant, and given by

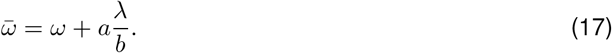

In Cartesian coordinates, along the limit cycle

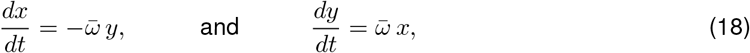

whose solution satisfying *x*(0) = 0 and *y*(0) = 1 is given by

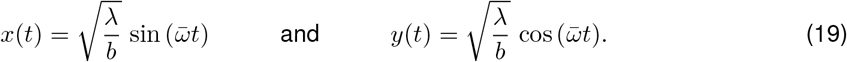

Solutions for other initial conditions are phase-shifts of this solution. This implies that limit cycles are identical and trajectories moving along the limit cycle have the same frequency. Fig. 4-A illustrates the *x*- and *y*-traces for representative parameter values. Fig. 4-B shows the corresponding phase-plane diagram, including the circular limit cycle characteristic of sinusoidal trajectories. Finally, Fig. 4-C shows the speed diagram, also characteristic of sinusoidal trajectories.

**Figure 4:**
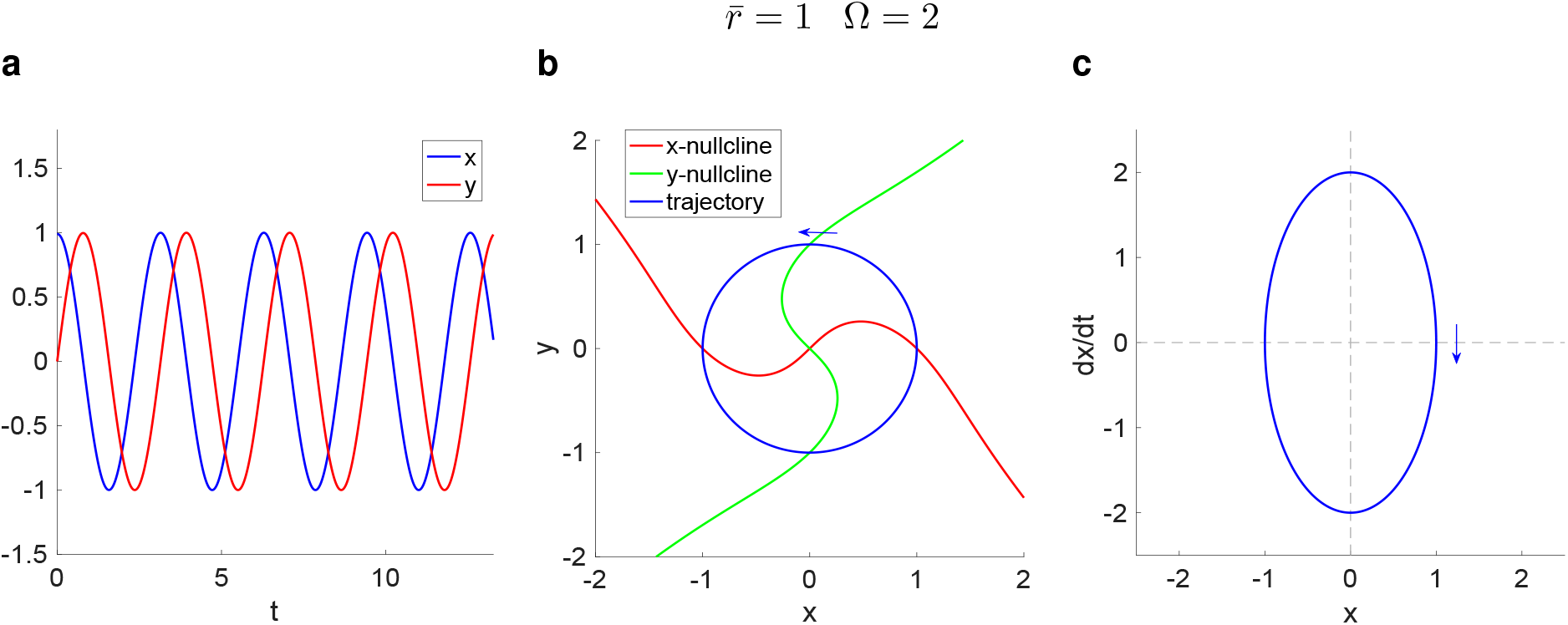
Dynamics of the *λ*-*ω* system for representative parameter values. **a.** Traces (curves of *x* and *y* as a function of *t*). **b.** Phase-plane diagram. The *x*- and *y*-nullclines are the set of state points in the *x*-*y* plane that make *dx/dt* = 0 and *dy/dt* = 0, respectively. **c.** Speed diagram for *x* (curves of *dx/dt* as a function of the state variable *x*). We used the following parameter values: *λ* = 1, *ω* = 1, *a* = 1 and *b* = 1.

#### Limit cycle degeneracy: the level sets for all possible attributes coincide

Along the limit cycle

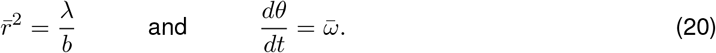

The limit cycles have the same amplitude for the infinitely many combinations of values *λ* and *b* for which their quotient is constant and they have the same frequency for the infinitely many combinations of the four parameters such that 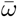 is constant. The amplitude level sets are curves in the two-dimensional *λ*-*b* parameter space and the frequency level sets are hypersurfaces in the four-dimensional *λ*-*b*-*ω*-*a* parameters space. Note that even along an amplitude level set there is frequency degeneracy along the limit cycle.

Fig. 5 illustrates the identical limit cycles for different combinations of parameter values. The different phase-plane diagrams supporting this limit cycle degeneracy indicate that the transient dynamics are different for the different cases. This is also clear from eq. (10). The time constant associated with the evolution of the envelope oscillations approaching the limit cycle (or moving away from it, if it is unstable) is *λ*-dependent.

**Figure 5:**
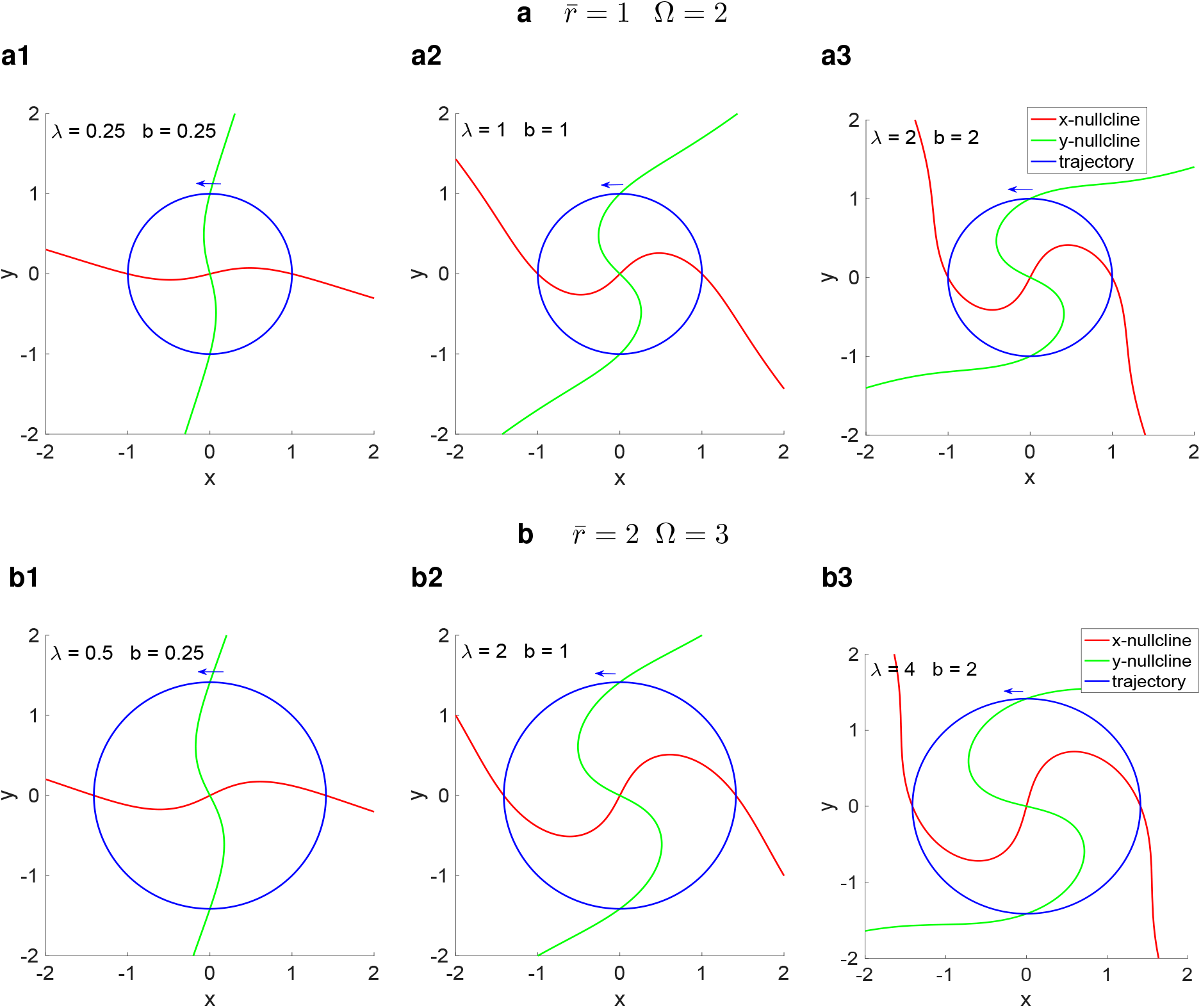
Phase-plane diagrams for the *λ*-*ω* system on two frequency and amplitude levels. **a** 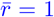 and Ω = 2. **b** 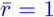 and Ω = 3. We used the following parameter values: *ω* = 1, *a* = 1.

**Figure 6:**
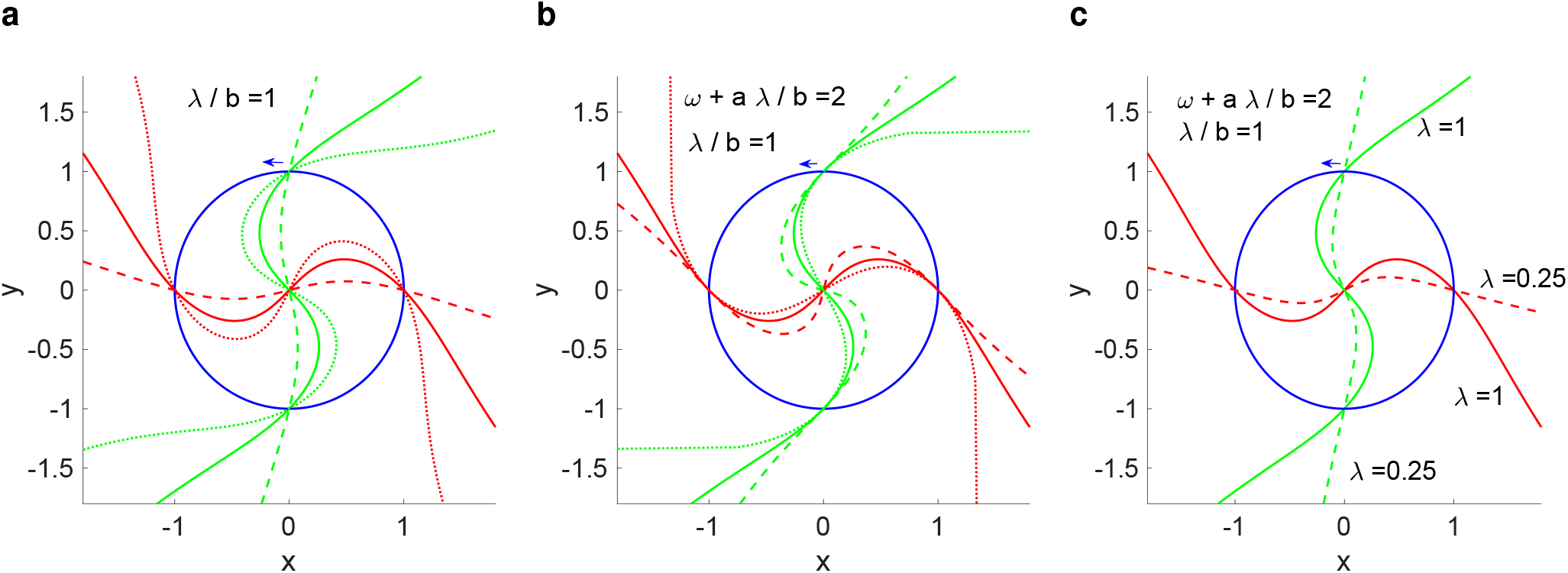
Compensation mechanisms for amplitude and frequency level sets in the *λ*-*ω*_2_ system. Each panel shows two superimposed phase-plane diagrams. The red curves represent the *x*-nullclines, the green curves represent the *y*-nullclines and the blue circles represent the limit cycles. a. Nullclines for values of *λ* and *b* on the same level set (*λ/b* = *const*) intersect at the same point lying on the limit cycle. We used the following parameter values: *λ* = *b* = 0.25 (dotted), *λ* = *b* = 1 (solid) and *λ* = *b* = 2 (dashed). We used *ω* = 1 and *a* = 1 for all cases. b. Nullclines for values of the four parameters on the same frequency level (*ω* + *a b/λ* = *const*) and values of *λ* and *b* on the same amplitude level set (*λ/b* = *const*) have the same slope at their point of intersection for the same value of *λ*. We used the following parameter values: *ω* = 1.75 and *a* = 0.25 (dotted), *ω* = 1 and *a* = 1 (solid) and *ω* = 0.25 and *a* = 1.75 (dotted). We used *λ* = 1 and *b* = 1 for all cases. c. The slope of the nullclines at their point of intersection depends on the values of *λ* for values of the four parameters on the same frequency level (*ω* + *a b/λ* = *const*) and values of *λ* and *b* on the same amplitude level set (*λ/b* = *const*). We used the following parameter values: *λ* = 1, *b* = 1, *ω* = 1 and *a* = 1 (solid) and *λ* = 0.25, *b* = 0.25, *ω* = 0.5 and *a* = 1.5 (dashed).

#### Limit cycle degeneracy: symmetries in the phase-plane diagram

The *x*- and *y*-nullclines for the *λ*-*ω* system are given by the solutions *y* = *F* (*x*) and *x* = *G*(*y*) of

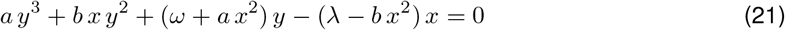

and

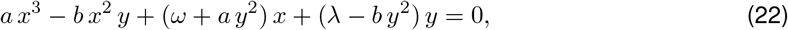

respectively.

The phase-plane diagram in Fig. 4-b shows the nullclines and trajectory for the solution to the *λ*-*ω* system for representative parameter values whose solution is shown in Fig. 4-a. We express the *y*nullcline as a function of *y*, and not of *x*, because as a function of *x* this nullcline is not invertible (Fig. 4-b).

System (14)–(15) is preserved under the following changes of coordinates: (i) (*x, y*) → (−*x, −y*), (ii) (*x, y*) → (−*y, x*), and (iii) (*x, y*) → (*y, −x*), but not under (*x, y*) → (*y, x*) and (*x, y*) → (−*y, −x*). The preservation property (i) reflects the fact that the two nullclines are odd-functions: *F* (*x*) = −*F* (−*x*) and *G*(*y*) = −*G*(−*y*). The preservation properties (ii) and (iii) reflect the fact that the two nullclines are the negative of one another *F* (*x*) = −*G*(*x*). Taken together, these properties reflect the fact that the two nullclines are orthogonally identical in the sense that they can be obtained from one another by a *π/*2 rotation (Fig. 4-b). The limit cycles are preserved if both nullclines are rotated by the same angle [74]. Fig. 5 illustrates that these asymmetries are preserved for parameter values for different values of the amplitude and frequency of the limit cycle.

### 3.4 Compensation mechanisms of generation of level sets in the λ-ω model

Here we investigate the dynamic mechanisms of generation of level sets in terms of the model parameters by focusing on the geometric properties of the phase-plane diagrams in Cartesian coordinates. We aim to identify the geometric constraints in the phase-plane diagrams for the parameter values that belong to the same level sets and give rise to the limit cycle degeneracy.

In our analysis, we assume to have no information about the properties of the limit cycles and level sets from the polar coordinates analysis above. The reason for doing so is to develop tools to analyze the models when the symmetries of the *λ*-*ω* model are broken and the level sets for the different attributes are split. In these cases Cartesian coordinates are more amenable for analysis than polar coordinates.

Because of the model symmetries reflected in the geometric properties of the *x*- and *y*-nullclines it is natural to assume that if a limit cycle exists, then it has circular shape and it is centered at the origin.

#### 3.4.1 Level sets in the *λ*-*b* parameter space

Setting *y* = 0 in the *x*-nullcline (21) shows that it crosses the *x*-axis at 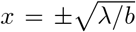, independently of the values of *ω* and *λ*. Likewise, setting *x* = 0 in the *y*-nullcline (22) shows that it crosses the *y*-axis at 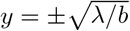, independently of the values of *ω* and *λ* (Fig. 7-a).

**Figure 7:**
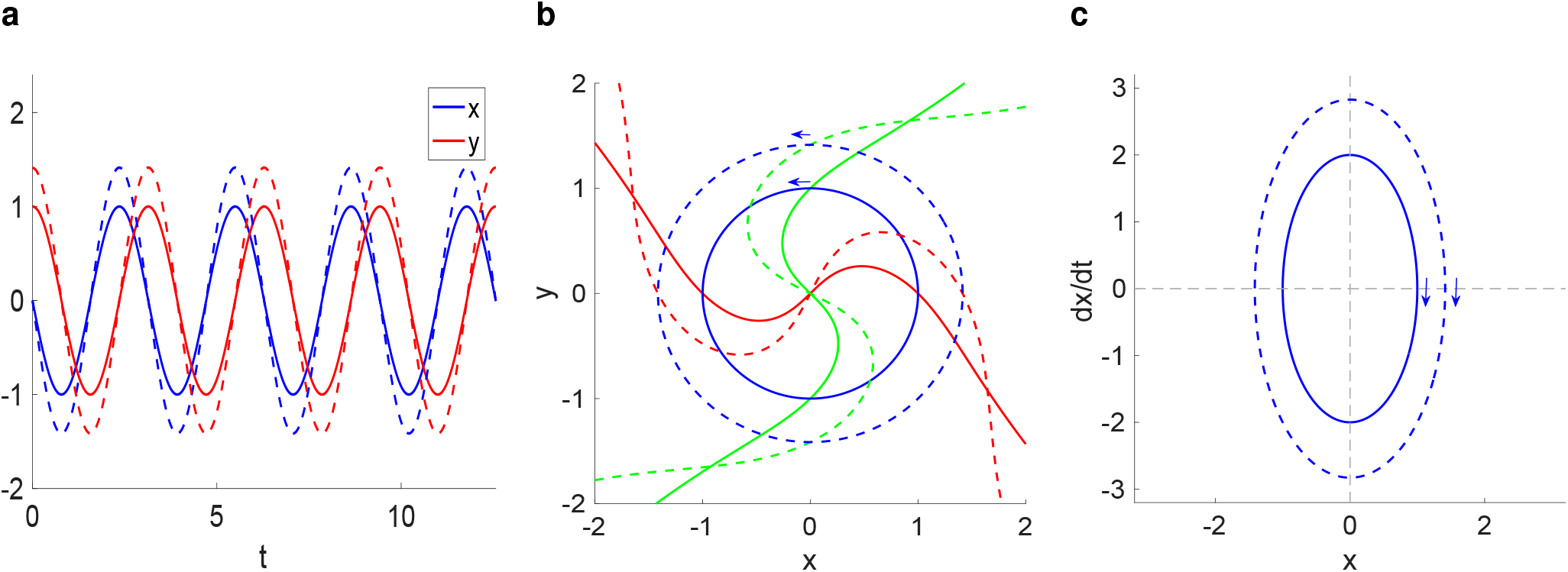
Dynamics compensation mechanisms in the *λ*-*ω*_2_ model for two sets of parameter values in the same frequency level set but different amplitude level sets. In all panels the frequency is *ω* + *a b/λ* = 2 and the amplitudes are 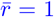 (solid) and 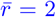 (dashed). **a.** Traces (*x*- and *y*-time courses). The dashed voltage traces evolve faster to compensate for the large limit cycle diameter. **b.** Superimposed phase-plane diagrams. The red curves represent the *x*-nullclines, the green curves represent the *y*-nullclines and the blue circles represent the limit cycles. **c.** Superimposed *x*-speed diagram. We used the following parameter values: *λ* = 1, *ω* = 1, *a* = 1 and *b* = 1 (solid) and *λ* = 2, *ω* = 1, *a* = 0.5 and *b* = 1.

The assumption that the limit cycle is circular and centered at the origin implies that the limit cycle trajectory is vertical on the *x*-axis and horizontal on the *y*-axis. This requires limit cycle trajectory to intersect the *x*-nullcline when the latter intersects the *x*-axis (at 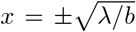) and to intersect the *y*-nullcline when the latter intersects the *y*-axis (at 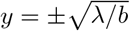). In other words, the intersection of the *x*-nullcline with the *x*-axis (or the intersection of the *y*-nullcline with the *y*-axis) determines the radius of the limit cycle trajectory without the need of the transformation to polar coordinates.

In general, changes in *λ* and *b* change the shape of the nullclines (compare Figs. 5-a, -b and -c). However, when *λ* and *b* change in such a way that *λ/b* remains constant, then the intersection of the *x*-nullcline with the *x*-axis is constant and therefore the radius of the limit cycle is constant. In other words, along a given level set *λ* and *b* compensate for each other so as to maintain the intersection of the *x*-nullcline with the *x*-axis constant (or, by symmetry, the intersection of the *y*-nullcline with the *y*-axis constant).

Along these level sets the frequency of the oscillations on the limit cycle is constant since from (14)–(15) with *λ* − *b r*^2^ = 0 it follows that

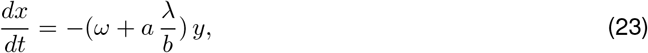

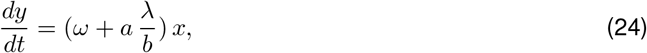

whose solution is a sinusoidal function of frequency *ω* + *a λb*^−1^ and phase tan^−1^(1). This implies that solutions belonging to the same level set are identical.

#### 3.4.2 frequency level sets in the *ω*-*a* parameter space: static compensation mechanisms when *λ* and *ω* belong to the same level set

Computation of the derivative of the *x*-nullcline (21) at 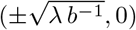 gives

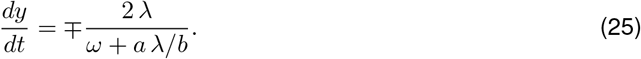

Similarly, computation of the derivative of the *y*-nullcline (22) at 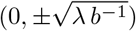 gives

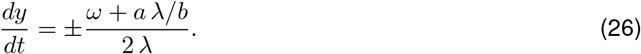

Therefore, the nullclines for parameter values on the same frequency level set (*ω* + *a b/λ* = *cont*) for the same value of *λ* are tangent at their point of intersection (Fig. 7-b). This constraint is lost when *λ* changes, even though the parameter values may belong to the same amplitude level set (Fig. 7-c).

#### 3.4.3 frequency level sets in the *ω*-*a* parameter space: dynamic compensation mechanisms when *λ* and *b* do not belong to the same level set

Here we focus on frequency level sets (*ω*+*a b/λ* = *const*) for parameter values such that *λ/b* ≠ *const*. In these cases the oscillation amplitudes are different (Fig. 8-a and -b). The *x*- and *y*-nullclines still intersect the corresponding limit cycles at the *x*- and *y*-axis, respectively, but, in order to maintain a constant frequency, the speed of the limit cycle trajectory needs to be greater than the limit cycle radius (Fig. 8-c).

**Figure 8:**
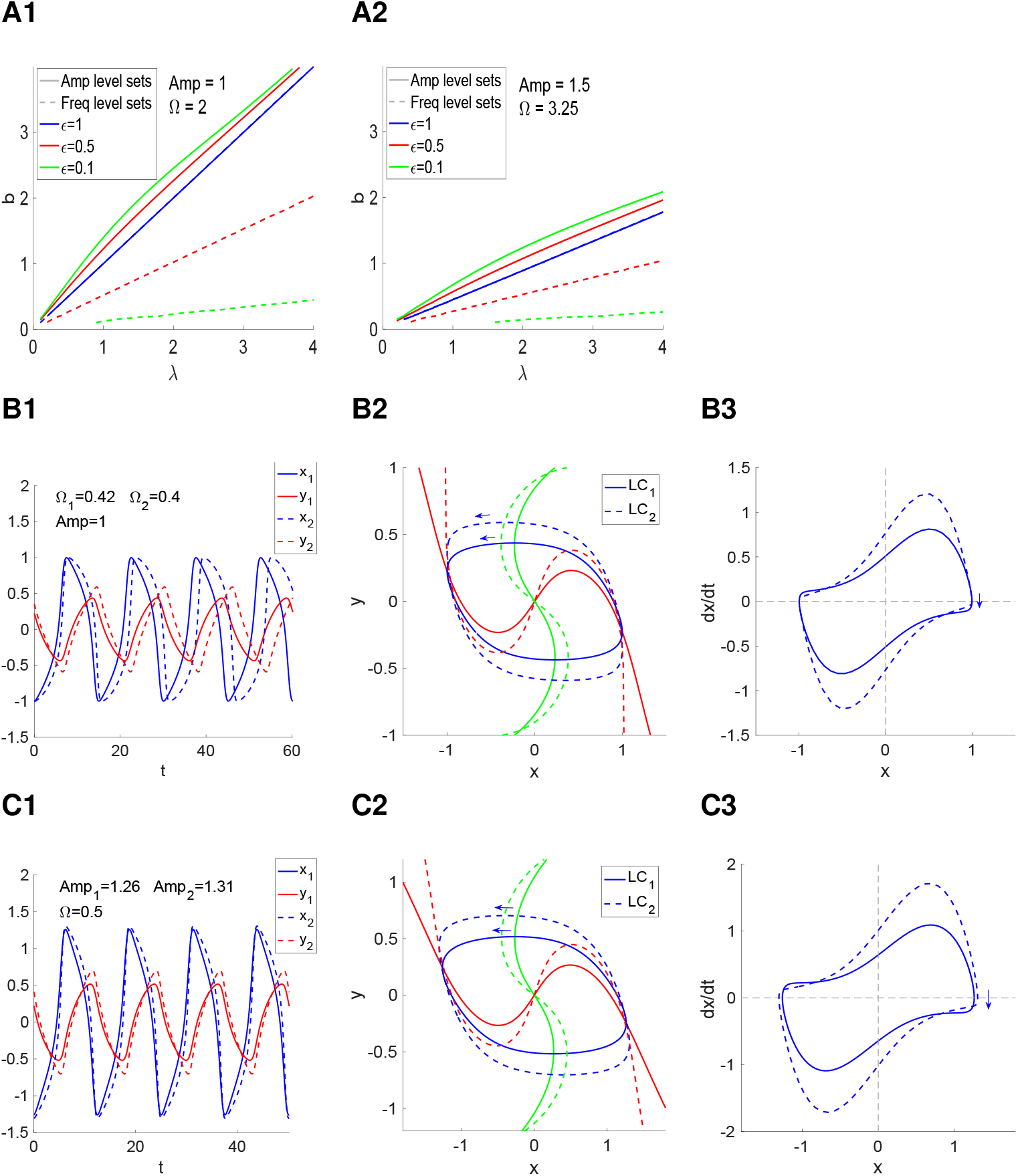
The canonical amplitude and frequency (Ω) level sets are not preserved under changes in the time scale separation (*ϵ*) between *x* and *y*. **A.** Amp (solid) and Freq (dashed) level sets in the *λ*-*b* parameter space for *ω* = *a* = 1 and representative values of *ϵ*. **A1.** *λ/b* = 1 and Ω = 2. **A2.** *λ/b* = 1.5 and Ω = 3.25. For *ϵ* = 1 (blue) the Amp and Freq level sets are superimposed. **B.** Representative examples of two oscillators on the same Amp (= 1) level set in A for *ϵ* = 0.1. **C.** Representative examples of two oscillators on the same Freq level set (Ω = 2) in A for *ϵ* = 0.1. **Left columns:** Superimposed traces for O_1_ and O_2_. **Middle columns:** Superimposed phase-plane diagrams for O_1_ and O_2_. The red curves represent the *x*-nullclines, the green curves represent the *y*-nullclines and the blue curves represent the limit cycles (LC_1_ and LC_2_) for the two oscillators. **Right columns:** Superimposed *x*-speed diagrams for O_1_ (solid) and O_2_ (dashed).

### 3.5 Unfolding of attribute level sets for different attributes due to symmetry breaking

The complete degeneracy of the limit cycle oscillations’ in the *λ*-*ω* model results from a number of symmetries present in the model, and a break in this symmetry is expected to disrupt the degeneracy. However, from previous work [32, 33] and our discussion above in the context of the HIV model (Fig. 2) we expect degeneracies to remain for the attributes characterizing the oscillation patterns. Here we address this issue by using the modified Λ-Ω model (7)–(8) where all the functions are quadratic and *r* is given by (6). The resulting modified *λ*-*ω* model reads

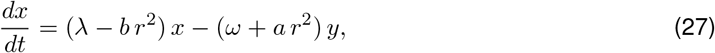

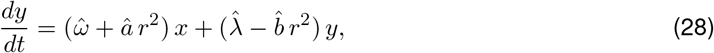

and reduces to the *λ*-*ω* model discussed above for 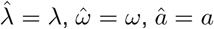 and 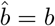.

Geometrically, from the dynamical systems point of view, there are four primary ways of breaking the symmetry between the *x*- and *y*-nullclines: (i) generating of a time scale separation between the two variables, (ii) expanding or shrinking the linear component of one equation with respect to the other, (iii) rotating one nullcline while keeping the other fixed, and (iv) displacing one nullcline while keeping the other fixed.

Here, we systematically analyze the effects of these symmetry breaking actions by choosing the appropriate values of the parameters in eq. (28) while keeping the parameter values in eq. (27) fixed. For simplicity, we focus on the amplitude and frequency level sets that emerge for each one of the models. In all cases, the canonical level sets are not preserved and new level sets emerge, which are different for different attributes and the four different ways of breaking the model symmetry. As expected, degeneracy remains for these attributes. As the parameter values move away from symmetry, both the shapes of the limit cycles and the shapes of the speed diagrams change, indicating that new dynamic compensation mechanisms to maintain the attributes’ degeneracy arise.

#### 3.5.1 Onset of frequency and amplitude level sets due to the presence of a time scale separation

A time scale separation between the first and second equations is generated by multiplying each one of the parameters in the *y*-equation by a parameter *ϵ*

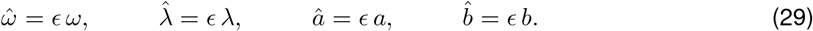

Fig. 8-A shows the frequency and amplitude level sets for representative parameter values. The presence of a time scale separation does not break the rotational symmetry of the two nullclines, but only the relative dynamics of the two variables, which is reflected in shapes of the limit cycles (Fig. 8-B2 and -C2) and the non-uniform speed along the limit cycle (Fig. 8-B3 and -C3) as well as the shapes of the oscillatory patterns (Fig. 8-B1 and -C1). The speed diagrams show the presence of dynamic compensation mechanisms responsible for the generation of the level sets.

#### 3.5.2 Onset of frequency and amplitude level sets due to changes in the linear properties of the *y*-nullcline

The region of linear behavior around the origin of the *y*-nullcline can be expanded or shrunk by multiplying the linear parameters *ω* and *λ* by a constant *c >* 1, while leaving the nonlinear parameters *a* and *b* unchanged

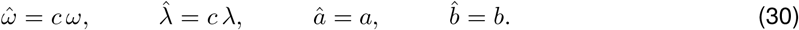

Fig. 9 shows our results for three values of *c*: *c* = 1 (symmetry), *c* = 1.25 and *c* = 1.5. The amplitude level sets almost coincide, but the frequency level sets are more separated (Fig. 9-A). This is reflected in the shapes of the limit cycles (Fig. 9-B2 and -C2) and speed diagrams ((Fig. 9-B3 and -C3). The speed diagrams show the presence of dynamic compensation mechanisms responsible for the generation of the level sets. These mechanisms are different from the ones discussed above.

**Figure 9:**
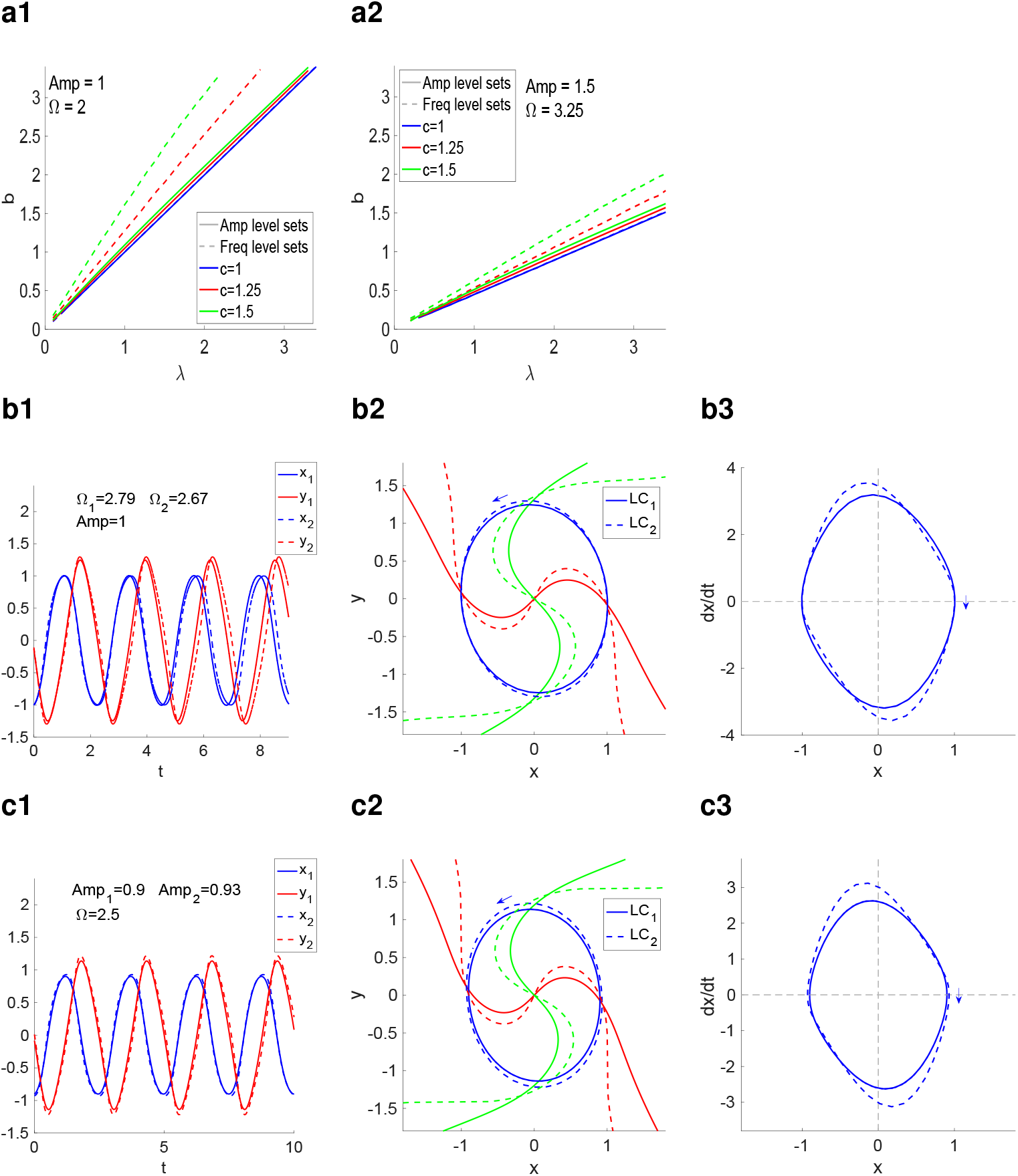
The canonical amplitude and frequency level sets are not preserved under changes in the linear properties of the *y*-nullcline. **A.** Amp (solid) and Freq (dashed) level sets in the *λ*-*b* parameter space for *ω* = *a* = 1 and representative values of the linearization coefficient *c*. **A1**. *λ/b* = 1 and Ω = 2. **A2.** *λ/b* = 1.5 and Ω = 3.25. For *ϵ* = 1 (blue) the Amp and Freq level sets are superimposed. **B.** Representative examples of two oscillators on the same Amp (= 1) level set in A for *c* = 2. **C.** Representative examples of two oscillators on the same Freq level set (Ω = 2) in A for *c* = 2. **Left columns:** Superimposed traces for O_1_ and O_2_. **Middle columns:** Superimposed phase-plane diagrams for O_1_ and O_2_. The red curves represent the *x*-nullclines, the green curves represent the *y*-nullclines and the blue curves represent the limit cycles (LC_1_ and LC_2_) for the two oscillators. **Right columns:** Superimposed *x*-speed diagrams for O_1_ (solid) and O_2_ (dashed).

#### 3.5.3 Onset of frequency and amplitude level sets due to rotation of the *y*-nullcline with respect to the *x*-nullcline

In order for the *y*-nullcline to be rotated by an angle *ϕ*, the parameters 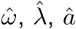 and 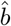 must be given by (see appendix B.1)

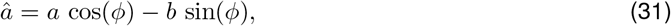

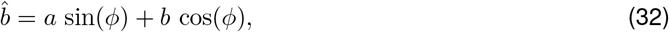

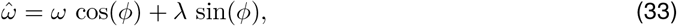

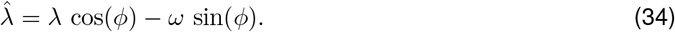

Fig. 10 shows our results for three values of *ϕ*: *ϕ* = 0 (symmetry), *ϕ* = *π/*6 and *ϕ* = *π/*3. The frequency and amplitude level sets do not coincide. The speed diagrams show the presence of dynamic compensation mechanisms responsible for the generation of the level sets. Comparison with the previous cases shows that these mechanisms are different from the ones discussed above. In particular, the speed diagrams in Figs. 10-B3 and -C3 have prominent intersections, which are almost entirely absent in the cases discussed above.

**Figure 10:**
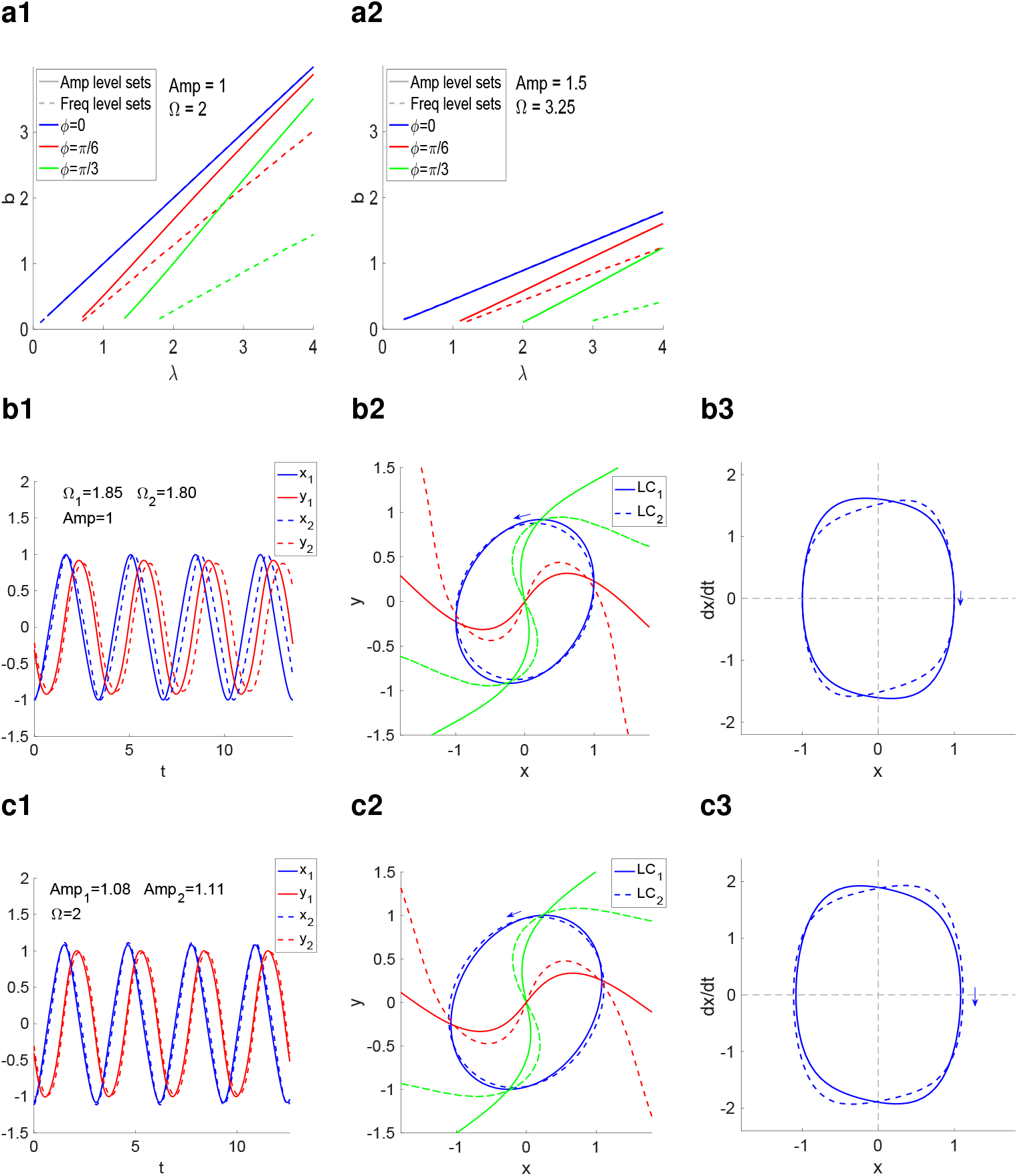
The canonical amplitude and frequency level sets are not preserved under rotations of the *y*-nullcline by an angle *ϕ* clockwise. **A.** Amp (solid) and Freq (dashed) level sets in the *λ*-*b* parameter space for *ω* = *a* = 1 and representative values of the rotation angle *ϕ*. **A1.** *λ/b* = 1 and Ω = 2. **A2.** *λ/b* = 1.5 and Ω = 3.25. For *ϕ* = 0 (blue) the Amp and Freq level sets are superimposed. **B.** Representative examples of two oscillators on the same Amp (= 1) level set in A for *ϕ* = *π/*6. **C.** Representative examples of two oscillators on the same Freq level set (Ω = 2) in A for *ϕ* = *π/*6. **Left columns:** Superimposed traces for O_1_ and O_2_. **Middle columns:** Superimposed phase-plane diagrams for O_1_ and O_2_. The red curves represent the *x*-nullclines, the green curves represent the *y*-nullclines and the blue curves represent the limit cycles (LC_1_ and LC_2_) for the two oscillators. **Right columns:** Superimposed *x*-speed diagrams for O_1_ (solid) and O_2_ (dashed).

#### 3.5.4 Onset of frequency and amplitude level sets due to the displacement of the *y*-nullcline with respect to the *x*-nullcline

The modified *λ*-*ω*_2_ system where the *y*-nullcline is displaced *x_c_* units in the horizontal direction from the position determined by system (14)–(15) is given by

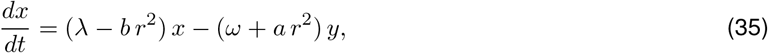

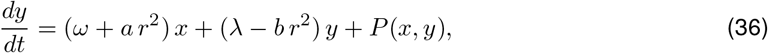

where

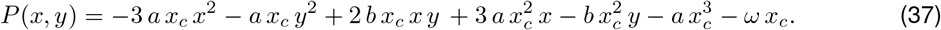

Fig. 11 shows our results for three values of *x_c_*: *x*_0_ = 0 (symmetry), *x_c_* = 0.2 and *x_c_* = 0.4. These results are similar to the previous cases, in that the frequency and amplitude level sets do not coincide and the speed diagrams show the presence of dynamic compensation mechanisms responsible for the generation of the level sets with prominent intersections (Figs. 11-B3 and -C3) as is only observed in Section 3.5.3.

**Figure 11:**
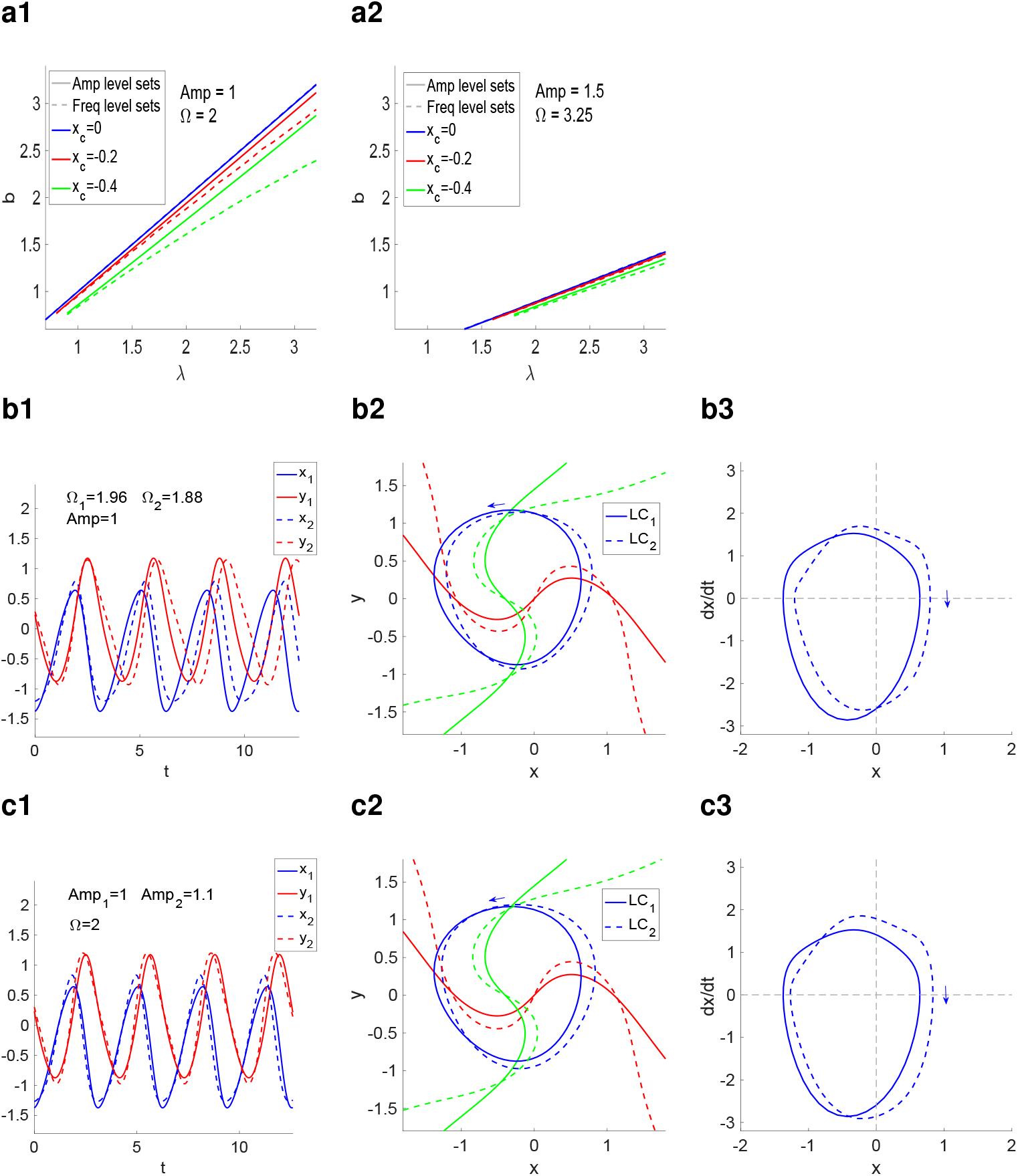
The canonical amplitude and frequency (Ω) level sets are not preserved under the displacement of the *y*-nullcline in a horizontal direction. **A.** Amp (solid) and Freq (dashed) level sets in the *λ*-*b* parameter space for *ω* = *a* = 1 and representative values of *x_c_*. **A1.** *λ/b* = 1 and Ω = 2. **A2.** *λ/b* = 1.5 and Ω = 3.25. For *ϵ* = 1 (blue) the Amp and Freq level sets are superimposed. **B.** Representative examples of two oscillators on the same Amp (= 1) level set in A for *x_c_* = −0.4. **C.** Representative examples of two oscillators on the same Freq level set (Ω = 2) in A for *x_c_* = −0.4. **Left columns:** Superimposed traces for O_1_ and O_2_. Middle columns: Superimposed phase-plane diagrams for O_1_ and O_2_. The red curves represent the *x*-nullclines, the green curves represent the *y*-nullclines and the blue curves represent the limit cycles (LC_1_ and LC_2_) for the two oscillators. **Right columns:** Superimposed *x*-speed diagrams for O_1_ (solid) and O_2_ (dashed).

### 3.6 Lambda-Omega systems of order 4: degeneracy, Hopf bifurcations and the emergence of bistability

Λ-Ω systems of order 4 (Λ-Ω_4_) correspond to the following quartic choices for Λ(*r*) and Ω(*r*)

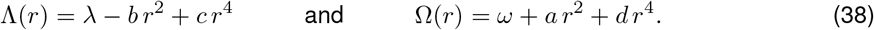

They are the normal forms of the Hopf bifurcation [84], and they have been used to model the dynamics of oscillatory systems in response to external inputs [85, 86].

The limit cycles, if they exist are given by

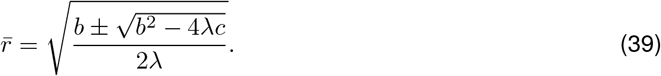

The fixed-points and limit cycles are stable (unstable) if 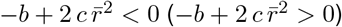.

Bistability emerges when there is a stable fixed-point, a stable limit cycle and an unstable limit cycle in between. There are a number of alternative scenarios that include the presence of one or two limit cycles. Along the limit cycles, the frequency is given by

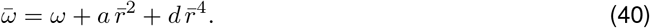

The degeneracies that emerge in this system, particularly in the bistable case, are more complex than in the *λ*-*ω* system discussed above and the level sets live in higher-dimensional parameter spaces. The analysis of these degeneracies and the compensation mechanisms that give rise to them goes along the same lines of the discussion above for the *λ*-*ω* system, however a detailed analysis is beyond the scope of this paper.

### 3.7 Degeneracy in extended λ-ω systems with generic waveforms: from sinusoidal to square and spike-like waveforms

The limit cycle in the standard *λ*-*ω* systems have sinusoidal waveforms. However, non-sinusoidal waveforms are ubiquitous in biological systems. Variations of the *λ*-*ω* system showing square-wave and spike-like waveforms have been introduced in [87]. Here we discuss the presence of degeneracy, in particular the complete limit cycle degeneracy in these systems.

The extended lambda-omega *λ*-*ω* systems (or extended Poincare oscillators) have the form

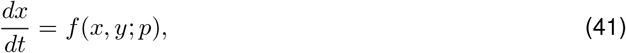

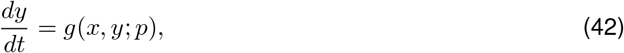

where

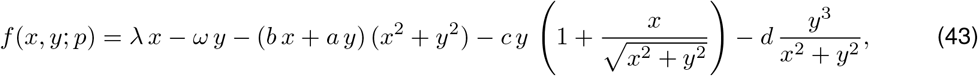

and

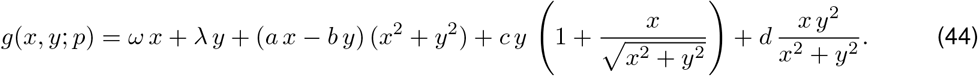

The parameters *p* (*ω*, *λ*, *a*, *b*, *c* and *d*) are non-negative constants. For *c* = *d* = 0, system (4)–(5) reduces to the standard *λ*-*ω* system. The parameters *λ* and *ω* control the linear dynamics, the parameters *a* and *b* control the cubic-like nonlinear terms (the nonlinearities in the standard *λ*-*ω* system) and the parameters *c* and *d* control the remaining nonlinear terms.

A change of coordinates from Cartesian to polar

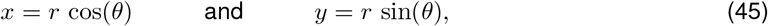

transforms system (4)–(5) into

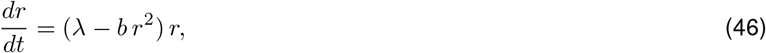

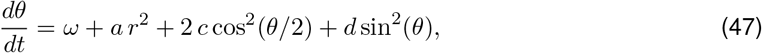

where *r* is given by (6)

#### Sinusoidal, spike-like and square-wave sustained oscillatory patterns for the extended *λ*-*ω* system

From eq. (10), the extended *λ*-*ω* system (4)–(5) has a circular limit cycle of radius

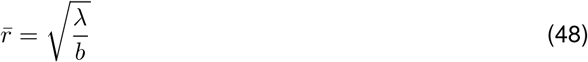

provided *λ/b >* 0 (Fig. 4). The limit cycle is stable if and only if *λ <* 0. Otherwise, the fixed-point 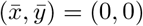 is stable. The properties of the right-hand side of eq. (73)

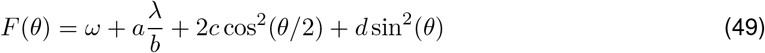

determine the various types of temporal patterns along these limit cycles.

For the *c* = *d* = 0, *F* (*θ*) is constant (Fig. 12-A2), and therefore *λ*-*ω* evolves linearly with time (Fig. 12-A3). The solutions are sinusoidal (Fig. 12-A1) with period

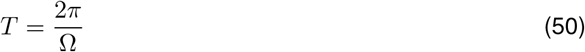

where

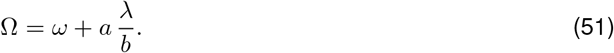

**Figure 12:**
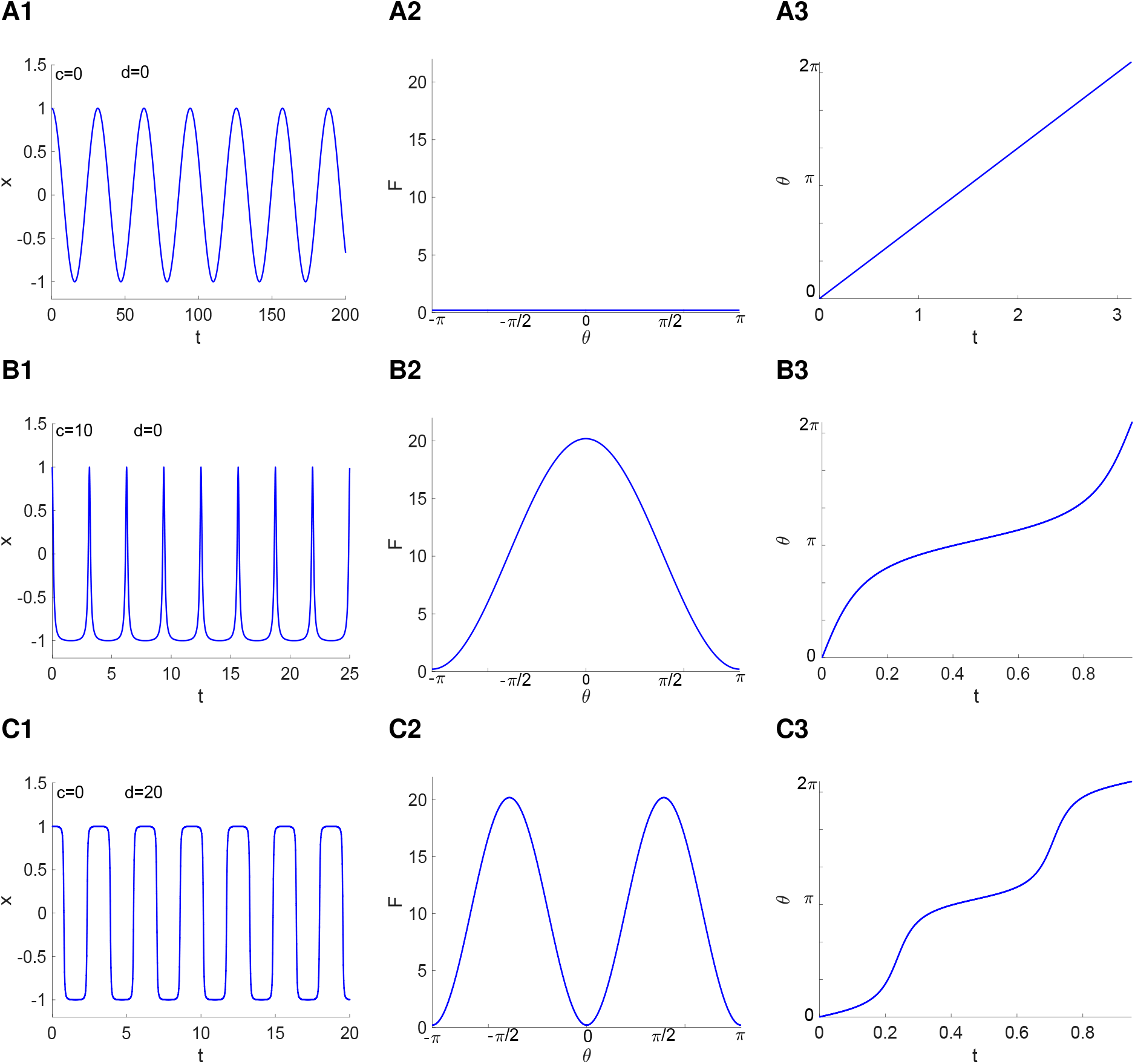
Representative limit cycle patterns within the same amplitude level set (*λ/b* = 1). The angular velocity is given by *F* (*θ*) = *ω* + *aλb*^−1^ + 2*c* cos^2^(*θ/*2) + *d* sin^2^(*θ*). **A.** Sinusoidal oscillations for uniform angular velocity (*c* = *d* = 0). **B.** Spike-like oscillations with slow-fast angular velocity (*c* = 10 and *d* = 0). *F* (*θ*) peaks once within the cycle at *θ* = 0. **C.** Square-wave oscillations with slow-fast angular velocity. *F* (*θ*) peaks twice within the cycle, at *θ* = −*π/*2 and *θ* = *π/*2. We used the following parameter values: *λ* = *b* = 1, *ω* = 0.1 and *a* = 0.1.

For *d* = 0, *F*(*θ*) peaks at *θ* = 0 (Fig. 12-B2) causing *θ* to evolve in a fast-slow manner (Fig. 12-B3) giving rise to spike-like patterns (Fig. 12-B1). The period is given by (see Appendix B)

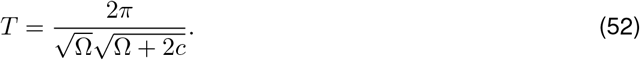

For *c* = 0, *F*(*θ*) peaks twice at *θ* = ±*π/*2 (Fig. 12-C2) causing *θ* to evolve in a fast slow manner, changing from fast to slow and vice versa twice within a cycle (Fig. 12-C3) and give rise to square-wave patterns (Fig. 12-C1). The period is given by (see Appendix B)

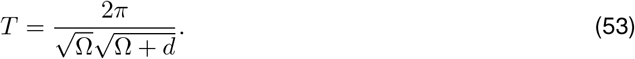

##### 3.7.1 Amplitude and period level sets

The amplitude level sets are as for the standard *λ*-*ω* system discussed above. The frequency level sets for the extended *λ*-*ω* system depend on the values of *c* and *d*. It is illustrative to consider two separate cases: (i) *c* = 0 and (ii) *d* = 0 for which the period *T* can be easily computed.

From (52), for *d* = 0, the period level sets are given by the solution to

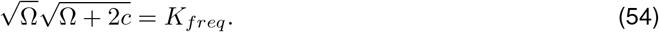

By solving one gets

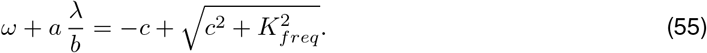

Similarly, from (52), for *c* = 0, the period level sets are given by the solution to

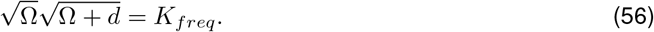

By solving one gets

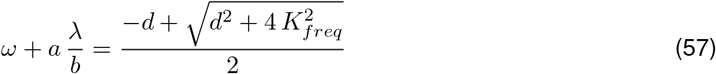

Similarly to the standard *λ*-*ω* system (*c* = *d* = 0), for all combinations of parameter values satisfying 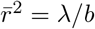 and either (55) or (57) the system has the same amplitude and frequency level sets.

#### 3.8 Parameter estimation algorithms recover at most parameter values on a level set

One potential way to disambiguate degenerate models is to obtain information about the transient behavior of trajectories. For example, for the *λ*-*ω* models, the parameter *λ* controls the evolution of the envelope of the oscillations with increasing amplitude converging to the limit cycle (with amplitude 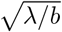). However, in realistic situations one does not have clean access to the transient evolution of trajectories unless one is able to perform perturbations to the oscillatory patterns. It has been argued that the presence of noise is useful to improve parameter estimation results since noise causes trajectories to explore wider regions of the phase-space and therefore more information about the dynamics, particularly the transient behavior of trajectories, is available to estimate parameters.

We used the two parameter estimation algorithms described in Section 2.4. In Fig. 13 we fixed the values of *ω* = *a* = 1 and attempted to estimate the values of *λ* and *b* using ground truth data (GTD) generated by using *λ* = *b* = *ω* = 1. In Fig. 14 we used the same GTD and attempted to estimate the four parameters.

**Figure 13:**
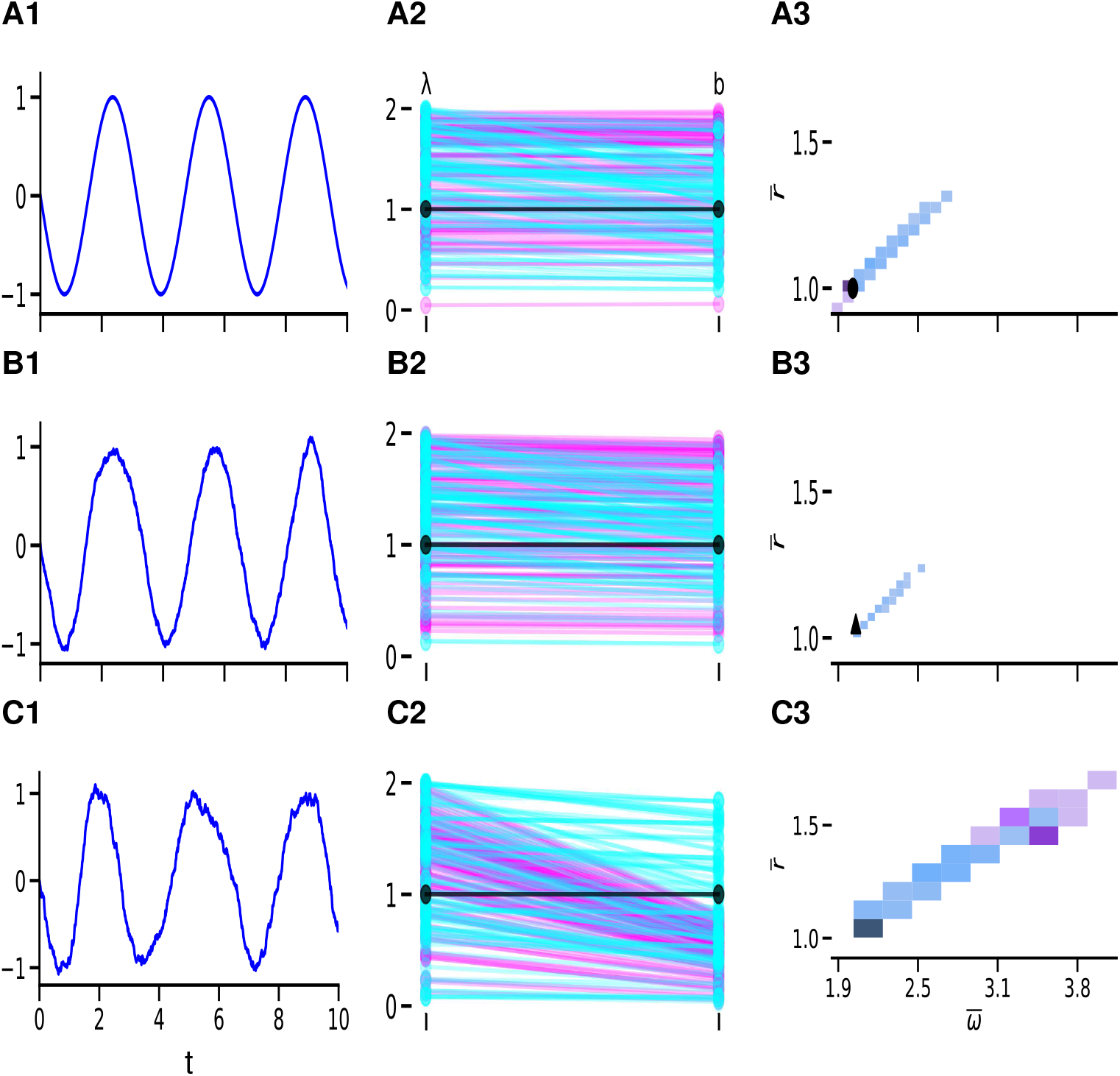
Parameter estimation algorithms recover at most parameter values on a level set. **Column 1.** Curves of *x* as a function of *t* for representative levels of noise increasing from top to bottom. **Column 2.** Recovered parameters (*λ* and *b*, for *ω* = *a* = 1) using a simulation based inference (SBI) approach (magenta) and genetic algorithm (GA) (cyan). The ground truth parameters used are *λ* = *b* = *ω* = *a* = 1 (black dots). Neither method is able to capture the true ground truth parameters. **Column 3.** Histogram of 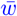 against 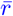 for the recovered parameter values (*λ* and *b*). Both parameter estimation approaches return parameter values that approximate the 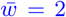 and 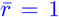 corresponding to the ground truth model parameters (black circles in column 2). Increasing levels of noise do not improve the estimates in either method. The attributes of the corresponding ground truth signal as measured by the peak detection routine (amplitude threshold = 0.1) are indicated by the circle, triangle, and square for respective levels of noise. Note the discrepancy between the values in panels C3 and A3. **A.** No noise. **B.** Intermediate level of noise. **C.** High level of noise.

**Figure 14:**
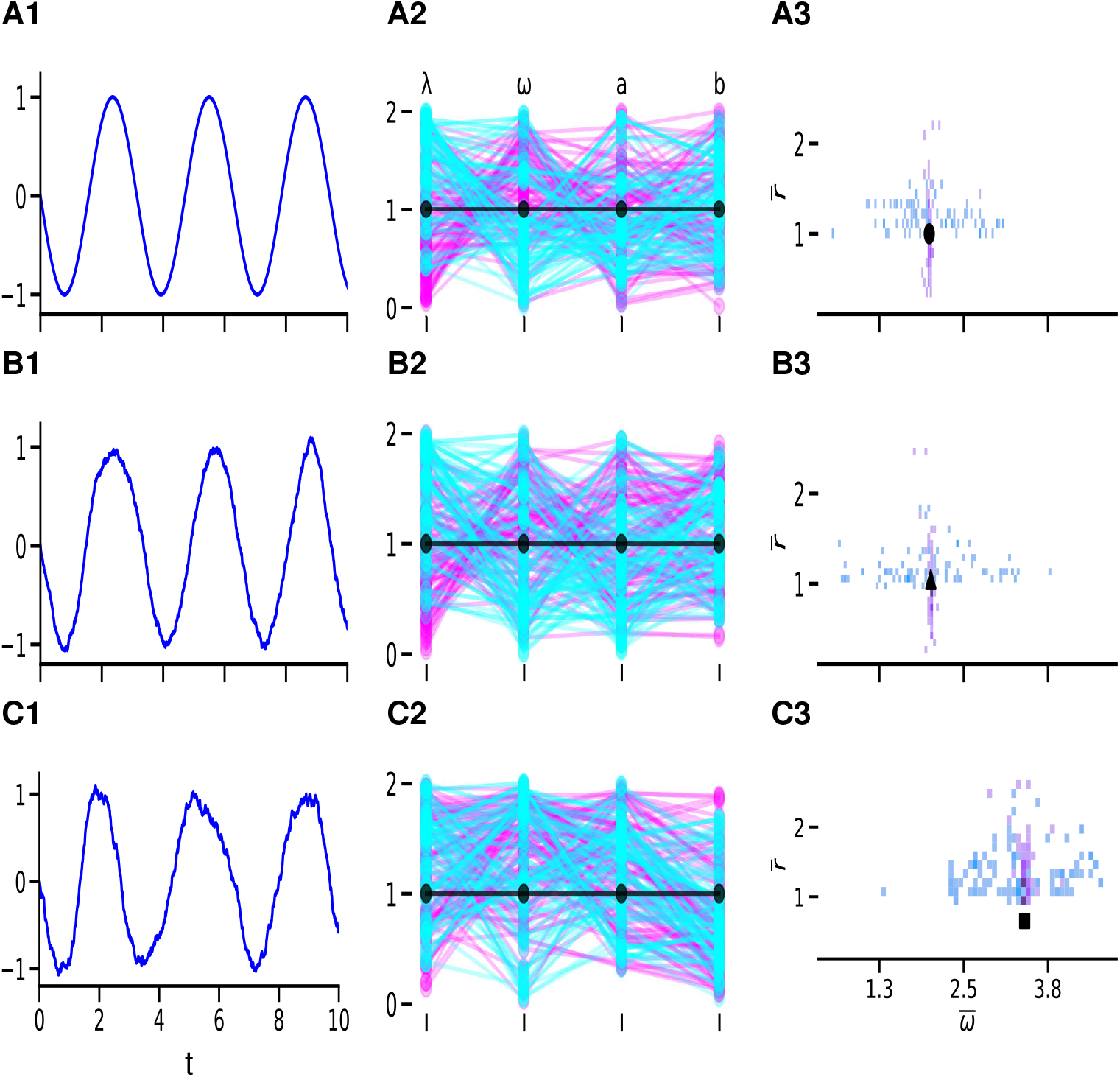
Parameter estimation algorithms recover at most parameter values on a level set. **Column 1.** Curves of *x* as a function of *t* for representative levels of noise increasing from top to bottom. **Column 2.** Recovered parameters (*λ*, *b*, *ω* and *a*) using a simulation based inference (SBI) approach (magenta) and genetic algorithm (GA) (cyan). The ground truth parameters used are *λ* = *b* = *ω* = *a* = 1 (black dots). Neither method is able to capture the true ground truth parameters. **Column 3.** Histogram of 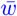 against 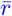 for the recovered parameter values (*λ*, *b*, *ω* and *a*). Both parameter estimation approaches return parameter values that approximate the 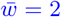 and 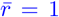 corresponding to the ground truth model parameters (black circles in column 2). Increasing levels of noise do not improve the estimates in either method. The attributes of the corresponding ground truth signal as measured by the peak detection routine (amplitude threshold = 0.1) are indicated by the circle, triangle, and square for respective levels of noise. Note the discrepancy between the values in panels C3 and A3. **A.** No noise. **B.** Intermediate level of noise. **C.** High level of noise.

Our results show that relatively small levels of noise are not enough to provide useful information, while higher levels of noise affect the efficacy of the parameter estimation algorithm. While these results are not conclusive, they illustrate that the degeneracy present in the models poses an obstacle to the ability to estimate the model parameters.

## 4 Discussion

Parameter estimation from experimental or observational data is a critical stage in any modeling study. There are several issues that conspire against the accurate estimation of model parameters, particularly the parameters in dynamic models (e.g, described by differential equations). Data are noisy and therefore one can at best expect to be able to identify a distribution of model parameters centered around the “true" parameter set (if it exists and is unique). In addition, one may not have access to data for all the state variables assumed to describe the process. Moreover, one may have experimental/observational access to discrete events (e.g., sequence neuronal spike times, record of a viral infection), but not the data that underlie the generation of these events (e.g., membrane potential, concentrations of the virus, infected and uninfected cells). Besides, if the data are too noisy or exhibit strong irregularities, one may only have reliable access to a number of activity attributes (e.g., oscillation preferred frequency, neuronal spiking mean firing rate) that may not be enough to capture the complete dynamic structure of these data (e.g., sinusoidal and relaxation oscillations may have the same frequency, amplitude and duty cycle). Furthermore, there may be gaps in the data sets and these gaps may be inconsistent across trials. Finally, even if one has clean experimental/observational access to all the state variables and the data is noiseless, the models may exhibit parameter degeneracy (multiple combinations of parameter give rise to the same pattern), rendering them structurally unidentifiable.

Classical structural unidentifiability is associated with the notion that one can at most identify combinations of unidentifiable model parameters [25,26]. This is due to an excess in the number of parameters with physical meaning as compared to the number of parameters necessary to compute the attributes that describe these data. We discussed these ideas in detail using a number of simple representative model examples with increasing levels of complexity in the Appendix A. A more thorough discussion can be found in [25,26] (and references therein).

In this paper we set out to analyze a different type of structural degeneracy/unidentifiability present in the family of Λ-Ω models [67–70, 87]. We found that Λ-Ω models have infinitely many parameter sets that produce identical stable oscillations, except possible for a phase-shift (reflecting the initial phase), but this parameters are not identifiable combinations of unidentifiable parameters as is the case in structural degeneracy. The number of model parameters in the Λ-Ω models is minimal in the sense that each one controls a different aspect of the model dynamics and the dynamic complexity of the system would be reduced by reducing the number of parameters. The level sets and the phase-plane diagrams can be computed analytically as well as the degenerate solutions. This adds clarity to the problem, leaving out other possible causes for unidentifiability (e.g., numerical, algorithmic), and facilitates the analytical investigation of the underlying compensation mechanisms.

Mathematically, degeneracy can be described in terms of level sets in parameter space for a given attribute (e.g., frequency, amplitude, duty cycle). An attribute level set is the set of all combinations of parameter values that produce solutions with a constant attribute [32]. In general, for a given model, level sets for different attributes may or may not coincide or be constrained by one another. For the Λ-Ω models, the level sets for all possible attributes coincide.

The family of Λ-Ω models produces a rich dynamic behavior, including monostable oscillations, bistability between oscillations and equilibria, Hopf bifurcations, and saddle-node bifurcations of limit cycles. The classical Λ-Ω models show sinusoidal oscillations, while the extended Λ-Ω models exhibit more complex waveforms, including square-wave (relaxation-type) and spike-like oscillations. Therefore, the Λ-Ω models serve as canonical models for degeneracy/unidentifiability for a large variety of realistic oscillatory processes. In addition, these canonical models can be used to investigate the consequences of degeneracy in single cells for their response to external perturbations [87] and the dynamics of networks in which these cells are embedded.

Degeneracy of oscillatory patterns has been the focus of various studies in the context of neuronal systems both experimentally and theoretically [32, 33, 64, 88–94]. Degeneracy is an inherent property of dynamical systems and is believed to arise from compensation mechanisms that generate balances among the participating processes (e.g., neuronal ionic currents) that control the dynamics [32,33]. Because the dynamic mechanisms of generation of biological oscillations involve the interplay of positive and negative feedback effects operating at different time scales, unidentifiability is likely to be present in the majority, if not all biological oscillatory systems. Therefore, the Λ-Ω models can serve as a first, guiding step to understand the mechanisms leading to degeneracy in biological oscillators and biological oscillatory networks [67,68].

The full degeneracy present in the Λ-Ω models results from the symmetries present in the model, which are captured by the phase-plane diagrams. We found that a systematic break of these symmetries for the *λ*-*ω* models leading to the phase-plane diagrams characteristic of more realistic models (e.g., relaxation oscillators) preserve the degeneracies of the attributes, although the level sets for different attributes do not necessarily coincide. Therefore, the *λ*-*ω* models serve as reference models to investigate the mechanisms of generation of attribute level sets for biological oscillators as perturbations to the corresponding *λ*-*ω* models that exhibits full degeneracy. We hypothesize that this is the case for the whole family of Λ-Ω models. More research is needed to test this hypothesis and clarify these issues.

In addition to biological oscillators, our results predict that the phenomenon of degeneracy may arise in other oscillatory systems in physics and chemistry since the *λ*-*ω* models are real-valued special cases of the complex Ginzburg-Landau equation [71, 72, 72–74, 95, 96]. Establishing these ideas requires additional research.

Unidentifiability in parameter estimation is a fundamental problem arising in the field of data science in connection to the models needed to analyze the available data. Degeneracy, the other side of the coin, is also a biological fact reported in biological experiments [64–66] (see also [32, 33]) and is expected to be pervasive in oscillatory systems. While on one hand degeneracy reflects the lack of information necessary to understand the underlying mechanism in term of the participating process, one the other hand, degeneracy has been proposed to endow systems with functional flexibility [97], thus adding a new dimension to the investigation of degenerate/unidentifiable systems. The family of Λ-Ω models serves as a framework to investigate these issues and is expected to have implications for the data-driven discovery of nonlinear dynamic equations [19,20]. Furthermore, the family of Λ-Ω models serves as the canonical models to calibrate parameter estimation algorithms. In contrast to the classical unidentifiable models, the family of Λ-Ω models is potentially identifiable if one has enough information about the transient behavior. However, as we pointed out above, this is information is not available in realistic situations. Our results using noisy ground truth data where transients are activated by noise were not useful to resolve the unidentifiability.

The family of Λ-Ω models also serves as a framework for the systematic investigation of degeneracy in dynamic models and the interplay between structural unidentifiability and the various forms of unidentifiability raised above resulting for lack of experimental/observational access to the state variables. The development of this framework and the implications for the investigation of degeneracy and identifiability in more realistic models requires more research.

## Acknowledgments

This work was partially supported by the National Science Foundation grants DMS-1715808 (HGR) and CRCNS-DMS-1608077, and a seed grant (HGR) awarded by the joint program between the New Jersey Institute of Technology (USA) and Ben Gurion University of the Negev (BGU, Israel). DL and RP were partially supported by NJIT Provost fellowships and NJIT Honors Summer Research Institute fellowships. We are grateful to Danny Barash and his research group (BGU) for useful discussions.

## A Model parameters, data-based activity attributes, unidentifiability and attribute level sets in simple models

## A.1 One-dimensional linear models

## A.1.1 Minimal model

We first consider the following 1D linear model

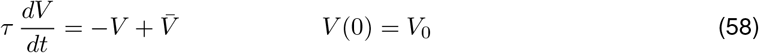

where *τ* is the time constant, 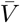 is the steady-state and *V*_0_ is the initial condition. The solution to eq. (58) is given by

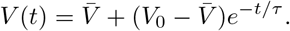

*V* decays (increases or decreases) monotonically towards the steady-state 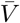 with a speed controlled by the time constant *τ* (Fig. 15-A). These parameters may or may not have any physical meaning, but they have a dynamic meaning in the sense that they are necessary (and sufficient) to describe the type of evolution curve shown in Fig. 15-A.

If the measured output is directly *V* (*t*), then *V*_0_ can be estimated from the initial conditions, 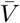 can be estimated from the steady state response lim_*t*→∞_ *V* (*t*), and *τ* can be estimated by computing the time it takes to *V* to reach 63 % of the gap between *V*_0_ and 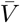 (Fig. 15-A). In this situation, all parameters are identifiable. They are structurally identifiable and remain practically identifiable when the available data is noisy, but a clean curve can be extracted by averaging across many trials, for example.

In this simple model scenario, the three model parameters are identical to the three activity *attributes* of the *V*-trace (curve of *V* as a function of *t*): the initial conditions, the time constant capturing the transient dynamics, and the steady-state (Fig. 15-A). We use the term attributes to refer to these parameters that characterize the data (observed data pattern). In these case, they are the minimal set of parameters that are necessary to characterize the data in terms of the model. If one can extract this information from the data, the error function is computationally less expensive by using the attributes instead of using the full data set.

If, instead, the measured output is *W*(*t*) = *kV*(*t*) where *k* is a (measurement) constant, then one can at most identify the three combinations of parameters *k V*_0_, 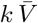 and *τ* by following the procedure described above, but not each one of the four parameters *V*_0_, 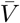, *τ* and *k*. Multiple combinations of values of *k* and *V*_0_ will satisfy *k V*_0_ = *W*_0_ = *W*(0) and multiple combinations of *k* and 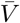 will satisfy 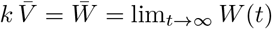.

Geometrically, this give rise to the *W*_0_ and 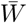 *level sets*, respectively, describing curves in the corresponding 2D parameter spaces (or surfaces/hypersurfaces in the 3D/higher-dimensional parameter spaces) for which *W*_0_ and 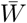 are constant. In the literature, this situation is referred to as the unidentifiable parameters (*V*_0_, *τ*, 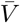 and *k*) forming identifiable parameter combinations (*k V*_0_, 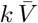 and *τ*) [25]. In other words, there is not enough information in the *V* traces to identify the four parameters. The number of model parameters has increased (four), but the number of attributes remains the same (three: *W*_0_, 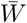 and *τ*).

## A.1.2 Model with realistic parameters

We consider here the following 1D linear model

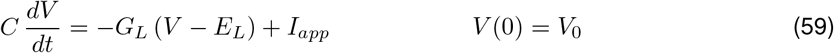

representing the dynamics of the membrane potential of a so-called passive cell, where *C* is the capacitance, *G*_*L*_ is the (passive or leak) conductance, *E*_*L*_ is the reversal potential and *I*_*app*_ is an applied (DC) constant current. This equation is obtained after applying Kirchhoff’s law to an electric circuit having a capacitor, a resistor and a DC input [98]. Each of the five parameters have a biophysical meaning.

By rescaling

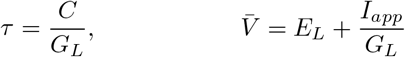

eq. (59) can be expressed in the form of eq. (58). These rescaled parameters have the dynamic meaning discussed above (they fully describe the evolution of the *V* -trace). If the measured output is *V*(*t*), then *V*_0_, 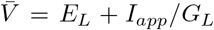 and *τ* = *C*/*G*_*L*_ can be estimated as for the minimal model discussed above. While the combinations *C*/*G*_*L*_, *E*_*L*_ + *I*_*app*_/*G*_*L*_ and *V*_0_ are identifiable, the set {*C*, *G*_*L*_, *E*_*L*_, *I*_*app*_, *V*_0_} is not and therefore the model is unidentifiable.

This (dynamic) redundancy in the biophysical parameters generate attribute level sets. The parameters *C* and *G*_*L*_ generate 2D *τ* -level sets and the parameters *E*_*L*_, *I*_*app*_ and *G*_*L*_ generate 3D 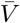 -level sets. Additional data and knowledge can come to the rescue. For example, if one knows *I*_*app*_ and one has data for two values of *I*_*app*_ (e.g., *I*_*app*_ = 0 and an additional non-zero value), then one can estimate *E*_*L*_ from 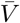 in response to *I*_*app*_ = 0, then one can estimate *G*_*L*_ form 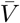 in response to *I*_*app*_ ≠ 0, by knowing this value, and then one can estimate *C* from *τ*. Following the same ideas as in Section A.1.1, the system remains unidentifiable (and more complex) if the output is *W*(*t*) = *kV* (*t*).

**Figure 15:**
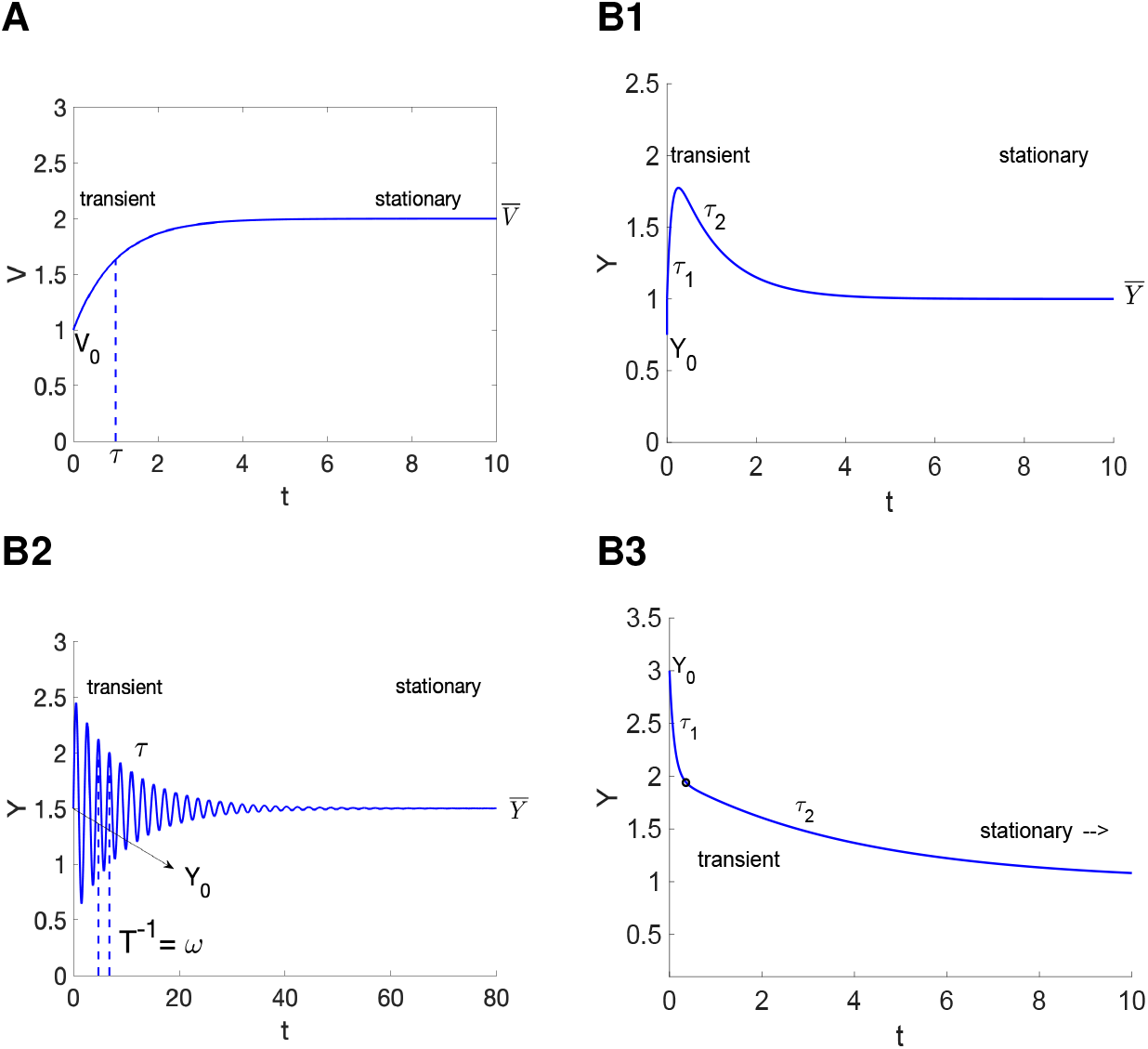
Activity attributes for simple models. **A.** 1D linear model. The model is defined by eq. (58). The activity attributes are the steady-state 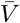, the time constant *τ* and the initial condition *V*_0_. The activity attributes coincide with the model parameters. **B.** 2D linear model. The model is defined by eq. (60). **B1.** Real eigenvalues (negative). *Y* is a difference of exponentials. The activity attributes are the steady-state 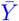, the time constants *τ*_1_ and *τ*_2_ (inverse of the eigenvalues), and the initial condition *Y*_0_. **B2.** Complex eigenvalues (negative real part). The activity attributes are the the steady-state 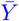, the frequency *ω* (equal to the inverse of the period *T*), the time constant *τ*, governing the evolution of the oscillations envelope, and the initial condition *Y*_0_. **B3.** Real eigenvalues (negative). *Y* is a sum of exponentials. The activity attributes are the steady-state 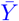, the time constants *τ*_1_ and *τ*_2_ (inverse of the eigenvalues), and the initial condition *Y*_0_. There is a separation of time scales: *τ*_1_ ≪ *τ*_2_.

## A.2 Two-dimensional linear models

## A.2.1 Minimal model

We consider the following 2D linear model

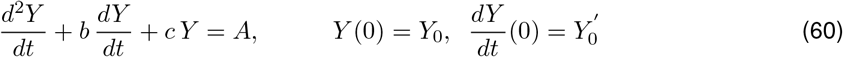

where *a*, *b* and *A* are constants. The steady state solution is given by

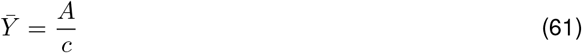

and the eigenvalues are given by

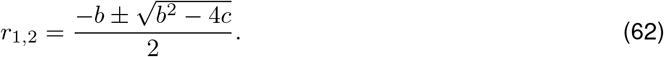

If the eigenvalues are real, the time constants are given by 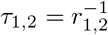. For certain parameter regimes, the solution exhibits an overshoot (Fig. 15-B1) or sag, which may not be detectable if one of the time constants is dominant, in which case the solution will look like a monotonically increasing (or decreasing) function. The attributes are the steady-state 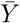, the time constants *τ*_1_ and *τ*_2_ and the initial condition *Y*_0_. The time constants could be replaced by other time metrics measuring the rise and decay phases of the solution during the transient overshoot. Alternative one of the time constants could be replaced by the overshoot peak (or sag trough). Note that the three of them are not independent attributes since one can be computed as a function of the others. For other parameter regimes, the solution is monotonically decreasing (Fig. 15-B3). When the time scales are well separated (e.g., *τ*_1_ ≪ *τ*_2_ in Fig. 15-B3) the solution is separated in two regimes: a rapid decay phase and a slower decay phase. The time constants for these two phases can be taken as attributes together with the steady state 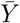. Alternatively, the approximate transition between the two (black dot n Fig. 15-B3) can be taken as an attribute in place of one of the time constants.

If the eigenvalues are complex, then the frequency of the damped oscillations is given by

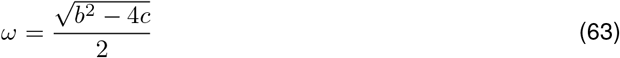

and the time constant governing the evolution of the damped oscillations amplitude is given by

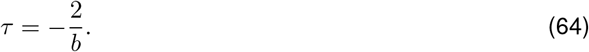

The solution exhibits damped oscillations (Fig. 15-B2). The attributes are the the steady-state 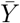, the frequency *ω* (equal to the inverse of the period *T*), the oscillation amplitude envelop time constant *τ* and the initial condition *Y*_0_.

In contrast to the 1D linear minimal model (58) discussed above, the attributes for the 2D linear minimal model (60) do not coincide with the model parameters. However, the former can be computed from the latter by appropriately using eqs. (61)–(64) and the initial conditions, and therefore they are identifiable. These (dynamic) parameters and the attributes they determine are necessary to reproduce the type of curves shown in Fig. 15-B.

As for the 1D linear minimal model, if the output is *Z*(*t*) = *k Y* (*t*) where *k* is a (measurement) constant, then the parameters *A* and *Y*_0_ become unidentifiable, while the products *kA* and *kY*_0_ are identifiable (combinations of unidentifiable parameters).

## A.2.2 Model with realistic parameters: two-node network

We consider here the following 2D linear system representing the dynamics of a simple network (Fig. 16) for the state variables *X*_1_ and *X*_2_

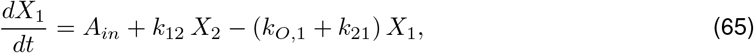

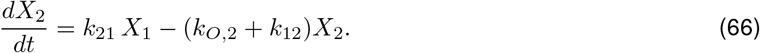

Cell 1 receives a constant input *A*_*in*_, interacts with cell 2 and the output is proportional to *X*_1_,

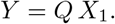

The other parameters are the transition constants (*k*_12_ and *k*_21_) for the interaction between cells 1 and 2, and the degradation (*k*_*O*1_ and *k*_*O*2_) constants for the two cells. The model has six unknown physical parameters (parameters with physical meaning). If one assumes the input *A*_*in*_ is known and the output is directly measured over *X*_1_, then the number of unknown parameters is reduced to four.

System (65)-(66) can be rewritten as a second order ODE in terms of the variable *Y*

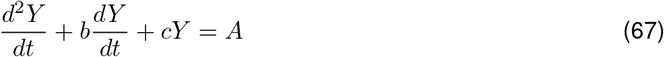

where

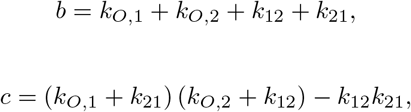

and

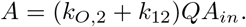

The parameters *b*, *c* and *A* are, in principle, identifiable following from our discussion about the activity attributes above, but the parameters *k*_*O*,1_, *k*_*O*,2_, *k*_12_, *k*_21_, *Q* and *A* are not. Even, by assuming *Q* = 1 (direct measurement of *X*_1_ and knowledge of the input *A*_*in*_, the remaining parameters are not identifiable. The physical parameters are dynamically redundant and create level sets in the corresponding parameters spaces.

**Figure 16:**
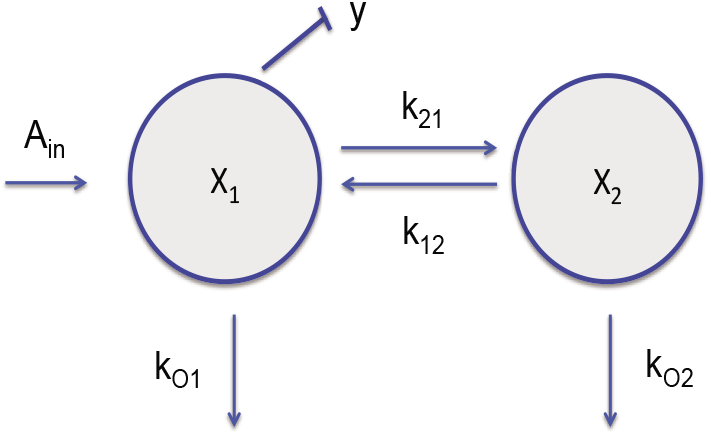
Diagram of a simple network.

## A.2.3 Model with realistic parameters: reduced hepatitis C virus (HCV) model during antiviral therapy

We consider here another 2D linear model [56] describing the dynamics of the infected cells *I* and the viral load *V*

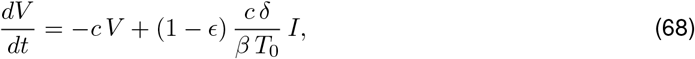

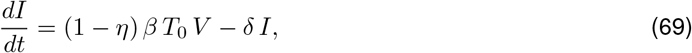

where *T*_0_ represent the target (uninfected) cells, assumed to be constant and equal to the target cell level at the beginning of the therapy, *β* represents the rate of infection, *δ* represents the rate of loss of the infected cell, *c* represents the rate of clearance of viral load. The rate of production of viral load is approximated by *cδ*/(*β T*_0_. This is the quasi-steady state of the 3D nonlinear model (for *T*, *V* and *I*) from which the linear model (68)–(69) was reduced [56] (see also [57]). The parameters *ϵ* and *η* represent the efficacy of the treatment in blocking the production of *V* and *I*, respectively. The initial conditions are *V*(0) = *V*_0_ and *I*(0) = *V*_0_*T*_0_*β/δ*, corresponding to the values of these variables at the steady state after therapy initiation [56].

System (68)–(69) can be rewritten as a second order ODE in terms of the variable *V*

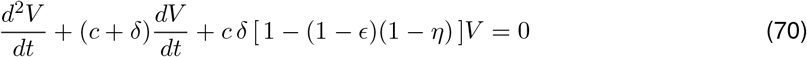

with *V*(0) = *V*_0_ and *dV*/*dt*(0) = −*cV*_0_*ϵ*. The coefficients of *dV/dt* and *V* in eq. (70) represent the parameters *b* and *c* in the minimal model (60).

For the parameter regimes considered in [56] (see also [57]), the solution decreases to *V* = 0 in two well separated phases, first very fast and then slower, similar to the evolution of *Y* in Fig. 15-B3. In this case, the two time constants *τ*_1_ and *τ*_2_ can be estimated.

The parameter *V*_0_ and the product *cϵ* can be computed from the initial conditions. The remaining parameters, either *c* or *ϵ*, *η* and *δ* need to be computed from the two attributes (the two eigenvalues or one eigenvalue and the transition point), therefore rendering the problem unidentifiable. We note that the unidentifiability could be stronger if one were not able to make some approximations (e.g, the initial condition *I*_0_).

## A.3 FitzHugh-Nagumo model: nonlinear oscillations

We use here the FitzHugh-Nagumo (FHN) model [49] in the following form

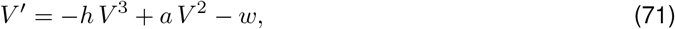

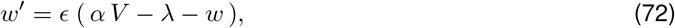

for the activation and inhibition variables *V* and *w*, respectively. In (71)–(72), *α*, *λ* and *ϵ* are constants, *ϵ* > 0, *α* > 0, and the *V* -nullcline *N*_*V*_ (*V*) = −*h V*^3^ + *a V*^2^ is a cubic function. Each one of these parameters plays a different dynamic role by controlling the geometry of the phase-plane diagram or representing the time constants. The parameters *h* and *a* control the shape of the *V* -nullcline in the phase-plane diagram. The parameters *α* and *λ* control the slope of the *w*-nullcline *N*_*w*_(*V*) = *αv* – λ and its position relative to the *V* -nullcline respectively. And the parameter *ϵ* represents the time scale separation between the two variables. For 0 < *ϵ* ≪ 1, the oscillations are of relaxation type, exhibiting abrupt transitions between the active and silent phases (Fig. 17-A and -B). As *ϵ* increases the oscillations, if they exist, transition from relaxation to oscillations of sinusoidal type. Fig. 17-C captures the two extremes of this type of transitions.

This model is minimal in the sense that each one of the parameters plays a dynamic role, and this parameter cannot be substituted by anther parameter, much in the same way was the cases discussed above. The FHN model has been used as a caricature model for various processes in nonlinear dynamics, including chemistry and neurobiology. Realistic models such as the Oregonator [99] (for the Belousov-Zhabotinsky chemical reaction [100, 101]) and the Morris-Lecar model [50, 51] (neuronal oscillations) have a similar dynamic structure to the FHN model in the sense that the nullclines in the phase-plane diagram are qualitatively similar.

The three natural attribute candidates are the oscillation frequency (or period), the oscillation amplitude and the duty cycle (fraction of the period that the variable is above its mean). However, these three attributes are not enough to differentiate between oscillations with different wave forms such as the ones presented in Fig. 17-C, and additional attributes are needed. One option is to incorporate the notion of a time constant capturing the time it takes for the variable to decrease from the peak to 37 % of the amplitude or the time it takes the variable to decrease from peak to trough.

**Figure 17:**
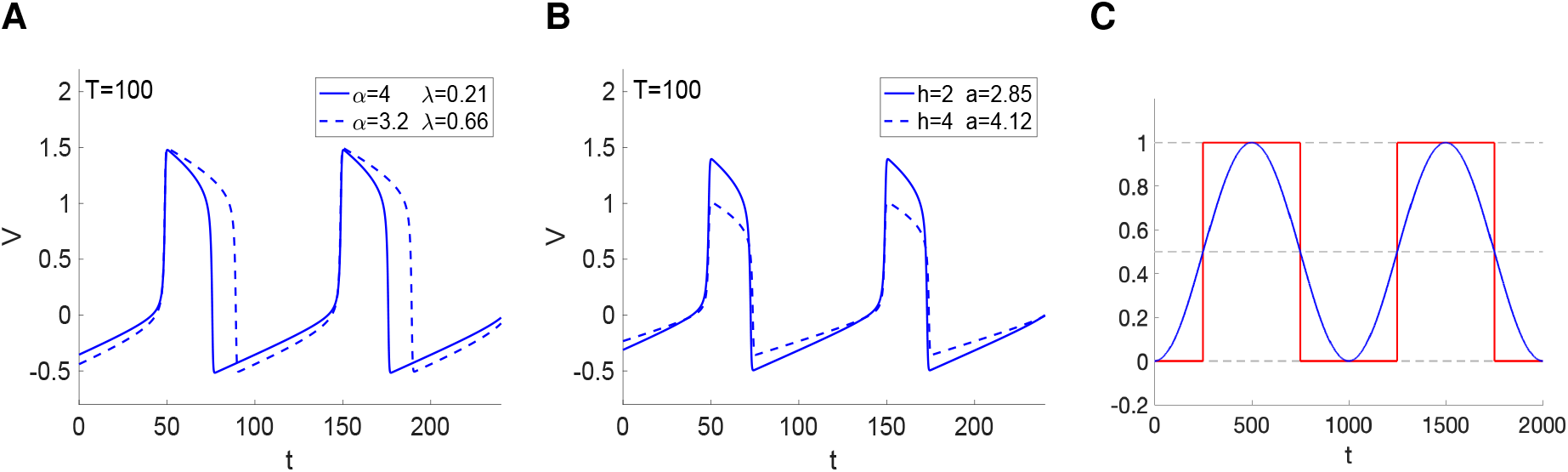
Period (frequency) level sets for the FHN model. **A.** Representative examples of oscillatory patterns belonging to a period (*T* = 100) level set in the (*α*,*λ*)-level set. We used *h* = 2, *a* = 3 and *ϵ* = 0.01. The two patterns belong to different duty-cycle level sets and are close to the same amplitude level set. **B.** Representative examples of oscillatory patterns belonging to a period (*T* = 100) level set in the (h,s)-level set. We used *α* = 4, *λ* = 0.1 and *ϵ* = 0.01. The two patterns belong to the same duty-cycle level set, but different amplitude level sets. **C.** Sinusoidal and square waves have the same period (frequency), amplitude and duty cycle level sets.

## B Solutions to the theta equation

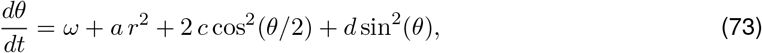

The solution to eq. (73) with *d* = 0 is given by

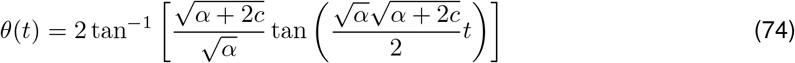

where

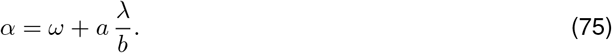

The solution to eq. (73) with *c* = 0 is given by

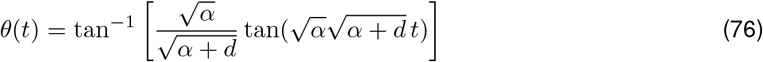

The computation of the period *T* in both cases requires taking into account the symmetric properties of the function *F*(*θ*). For the first integral it is enough to compute twice the integral in [0, *π*] and for the second integral it is enough to compute four times the integral in [0, *π*/2].

## B.1 Rotated lambda-omega systems of order 2

Each point in the *x*- and *y*-nullclines (16)–(21) can be expressed in polar form as

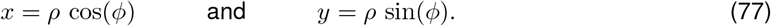

Substitution into (16)–(21) yields equations for the *x*- and *y*-nullclines in polar form

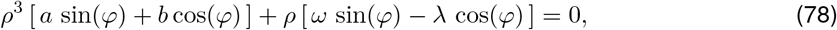

and

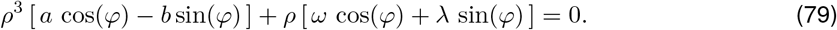

In order to rotate the nullclines by an angle *α* while preserving their shape we substitute *ϕ* by *ϕ* + *α* in (78)–(79), rearrange terms and define

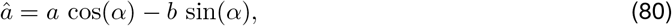

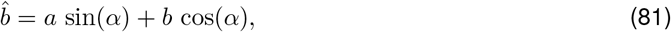

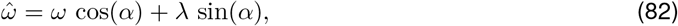

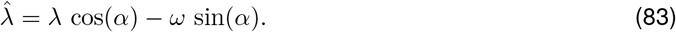

The resulting nullclines have the form (78)–(79) with *ω*, *λ*, *a* and *b* substituted by 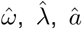 and 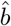, respectively.

In Cartesian coordinates, the resulting rotated *λ*-*ω*_2_ system is given by

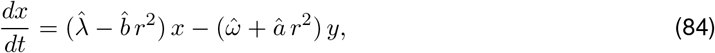

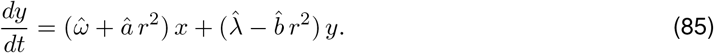

